# Universal recording of cell-cell contacts *in vivo* for interaction-based transcriptomics

**DOI:** 10.1101/2023.03.16.533003

**Authors:** Sandra Nakandakari-Higa, Maria C. C. Canesso, Sarah Walker, Aleksey Chudnovskiy, Johanne T. Jacobsen, Jana Bilanovic, S. Martina Parigi, Karol Fiedorczuk, Elaine Fuchs, Angelina M. Bilate, Giulia Pasqual, Daniel Mucida, Yuri Pritykin, Gabriel D. Victora

**Author notes:** These authors contributed equally (Maria C. C. Canesso, Sarah Walker).

## Abstract

Cellular interactions are essential for tissue organization and functionality. In particular, immune cells rely on direct and usually transient interactions with other immune and non-immune populations to specify and regulate their function. To study these “kiss-and-run” interactions directly *in vivo*, we previously developed LIPSTIC (Labeling Immune Partnerships by SorTagging Intercellular Contacts), an approach that uses enzymatic transfer of a labeled substrate between the molecular partners CD40L and CD40 to label interacting cells. Reliance on this pathway limited the use of LIPSTIC to measuring interactions between CD4^+^ helper T cells and antigen presenting cells, however. Here, we report the development of a universal version of LIPSTIC (uLIPSTIC), which can record physical interactions both among immune cells and between immune and non-immune populations irrespective of the receptors and ligands involved. We show that uLIPSTIC can be used, among other things, to monitor the priming of CD8^+^ T cells by dendritic cells, reveal the cellular partners of regulatory T cells in steady state, and identify germinal center (GC)-resident T follicular helper (Tfh) cells based on their ability to interact cognately with GC B cells. By coupling uLIPSTIC with single-cell transcriptomics, we build a catalog of the immune populations that physically interact with intestinal epithelial cells (IECs) and find evidence of stepwise acquisition of the ability to interact with IECs as CD4^+^ T cells adapt to residence in the intestinal tissue. Thus, uLIPSTIC provides a broadly useful technology for measuring and understanding cell–cell interactions across multiple biological systems.

## Introduction

Physical interactions in which cells use membrane-bound molecules to exchange signals with each other are at the core of multiple tissue functions^1, 2^. In the immune system they feature prominently, from the priming of T cells by dendritic cells (DCs) that initiates the adaptive immune response to the CD4^+^ T cell that enables B cells to produce high-affinity antibodies^3, 4^. More recent work has explored the role of similar interactions between immune and non-immune cells, such as those forming the epithelial barrier of the gut and skin, which are thought to drive transcriptional changes in immune cells that in turn enable them to support tissue function^5–8^. Despite their importance, direct observation of cell-cell interactions has traditionally been done by microscopy and other forms of imaging^9, 10^, which have the common limitation that interacting cells cannot be retrieved for downstream analysis^11^. Thus, the impact of the interaction on cell behavior and the cellular features that lead the interaction to occur in the first place cannot be inferred from traditional imaging data alone. More recently, spatial transcriptomics and high-density imaging technologies have allowed for a more in-depth characterization of the states of cells in the same neighborhood^12^. Although the more widely available technologies involve a tradeoff between high spatial resolution^13^ and full transcriptome coverage^14^, custom platforms can achieve gene expression profiling at cellular size resolution^15, 16^. However, even when capable of high resolution, transcriptomic and imaging techniques still report on proximity between cells rather than on true physical interaction and signal exchange between membranes, requiring additional indirect methods and assumptions to infer functional interactions computationally^17–22^. High throughput identification of cellular interactors and full deconvolution of the transcriptomic effects of physical interaction on cellular behavior and function are therefore yet to be achieved.

An attractive strategy to overcome these limitations has been proximity-based labeling across cellular membranes^23–27^. These strategies rely on equipping “donor” cells with enzymes or other signals that act over short distances to identify “acceptor” cells in either close proximity or physical contact with these donors. An early example of such an approach was our development of LIPSTIC, which uses enzymatic labeling across immune synapses to directly record cell-cell interactions *in vivo^24^*. In its first iteration, we engineered LIPSTIC to label interactions driven by a specific receptor-ligand pair—CD40L-CD40—a dependency that restricted its utility to interactions involving effector CD4^+^ T cells, the primary expressors of CD40L. Here, we report the development of a universal (u)LIPSTIC tool, which enables us to record interactions between multiple cell types of immune and non-immune origin. Coupling of uLIPSTIC with standard single-cell mRNA sequencing (scRNA-seq) methods allows for atlas-type characterization of the “cellular interactome” of a population of interest and for the definition of the molecular pathways associated with such interactions. Thus, uLIPSTIC enables us to achieve truly quantitative interaction-based transcriptomics without need for computational inference of transcriptomes or interacting molecules.

## Results

### A receptor/ligand-agnostic approach for recording cell–cell interactions

LIPSTIC uses the *Staphylococcus aureus* transpeptidase Sortase A (SrtA)^28^ to covalently transfer a peptide substrate containing the motif LPETG onto an N-terminal pentaglycine (G_5_) acceptor. In its original version, catalysis by the very low-affinity (∼1.8 mM) interaction between LPXTG-loaded SrtA and its G_5_ target^29, 30^ was favored by genetically fusing each component to one of the members of a receptor–ligand pair^24^, thus raising local concentration of the reactants above the threshold required for substrate transfer (**Fig. 1A**). We reasoned that a similarly high local concentration of enzyme and target could also be achieved in a receptor–ligand independent manner by driving very high expression of SrtA and G_5_ on apposing cell membranes without direct fusion to the interacting molecules. Transfer of substrate under these conditions would provide a universal readout for physical interactions between cells of any type, even in the absence of prior knowledge of the surface molecules that drive these interactions (**Fig. 1B**).

To test this strategy, we generated a donor–acceptor pair consisting of the “PDK” version of SrtA^30^ targeted to the plasma membrane by fusion to the human PDGFRB transmembrane domain^24^ (mSrtA) and the G_5_ peptide fused to the N-terminus of the mouse Thy1.1 protein, a single glycosyl-phosphatidyl-inositol (GPI)-anchored immunoglobulin domain which is widely used as both an allelic marker and a reporter for retroviral transduction. 3D modeling (**Fig. 1C**) predicted the maximal distance between membranes at which label transfer would occur to be approximately 14 nm (the total distance spanned when the PDGFRB N-terminal random coil is fully extended). This distance is comparable to the inter-membrane span required, for example, for the TCR-MHC interaction (∼15 nm), and narrower than the typical distance separating juxtaposed cell membranes in the absence of receptor-ligand interactions, set by glycocalyx repulsion^31^. Given the negligible affinity (∼1.8 mM) between SrtA-LPETG and G5 ^30^, such a design would in principle allow for label transfer only when cells were functionally interacting at a close intermembrane distance, without driving artificial interactions between its engineered components.

To gauge the functionality of such an approach, we first transfected separate pools of HEK293T cells with high or low concentrations of plasmids expressing either mSrtA or G_5_-Thy1.1, then carried out uLIPSTIC labeling by addition of biotin-LPETG substrate to co-cultures of these cells as described^24^ (**Fig. 1D**). Labeling of G_5_-Thy1.1-transfected cells was augmented when donor and acceptor populations were forced to physically interact by co-transfection of constructs encoding CD40L and CD40, respectively, and further increased when the uLIPSTIC components were transfected at the highest concentration (**Fig. 1E**). Thus, high-level expression of SrtA and G_5_ on the membranes of interacting cells allows for LIPSTIC labeling to proceed without need for fusion to specific receptor–ligand pairs, in principle enabling universal labeling of physical interactions between cells of any type.

Based on these initial findings, we generated a *Rosa26*^uLIPSTIC^ mouse allele in which high expression of mSrtA (preceded by a FLAG-tag) or G_5_-Thy1.1 are driven by the strong cytomegalovirus early enhancer/chicken beta-actin/rabbit beta-globin (CAG) promoter introduced into the ubiquitously expressed *Rosa26* locus, using a previously published targeting strategy^32^ (**Fig. 1F** and **Fig. S1**). The upstream G_5_-Thy1.1 is flanked by LoxP sites, so that Cre-mediated recombination leads to expression of a previously silent downstream mSrtA, switching Cre-expressing cells from uLIPSTIC acceptors into uLIPSTIC donors. The specificity and efficiency of this recombination are determined by the Cre driver used (**Fig. S2**). To test this system in primary immune cells *ex vivo*, we crossed *Rosa26*^uLIPSTIC^ mice to the CD4-Cre and OT-II T cell receptor transgenes to generate mSrtA^+^ uLIPSTIC donor T cells specific for peptide 323-339 of the model antigen chicken ovalbumin (OVA). Co-culture of mSrtA^+^ donor T cells with G_5_-Thy1.1^+^ B cells showed efficient transfer of labeled substrate between T and B cells only in the presence of OVA_323-339_, whereas background labeling by non-interacting cells in the absence of OVA_323-339_ was negligible (**Fig. 1G**). Substrate transfer was completely prevented by addition of a blocking antibody to MHC-II, necessary for the cognate recognition of B cells by T cells, but not by an antibody to CD40L, which is not itself required to drive this interaction (**Fig. 1G**). We conclude that uLIPSTIC provides an approach for trans-synaptic labeling of intercellular contacts that is functional regardless which receptor(s) and ligand(s) drive these interactions. Importantly, preventing functional B cell–T cell interactions by blocking antigen presentation completely abrogated label transfer between cells, indicating that high expression of the uLIPSTIC components is itself not sufficient to drive unwarranted cell–cell interactions.

**Figure 1.**
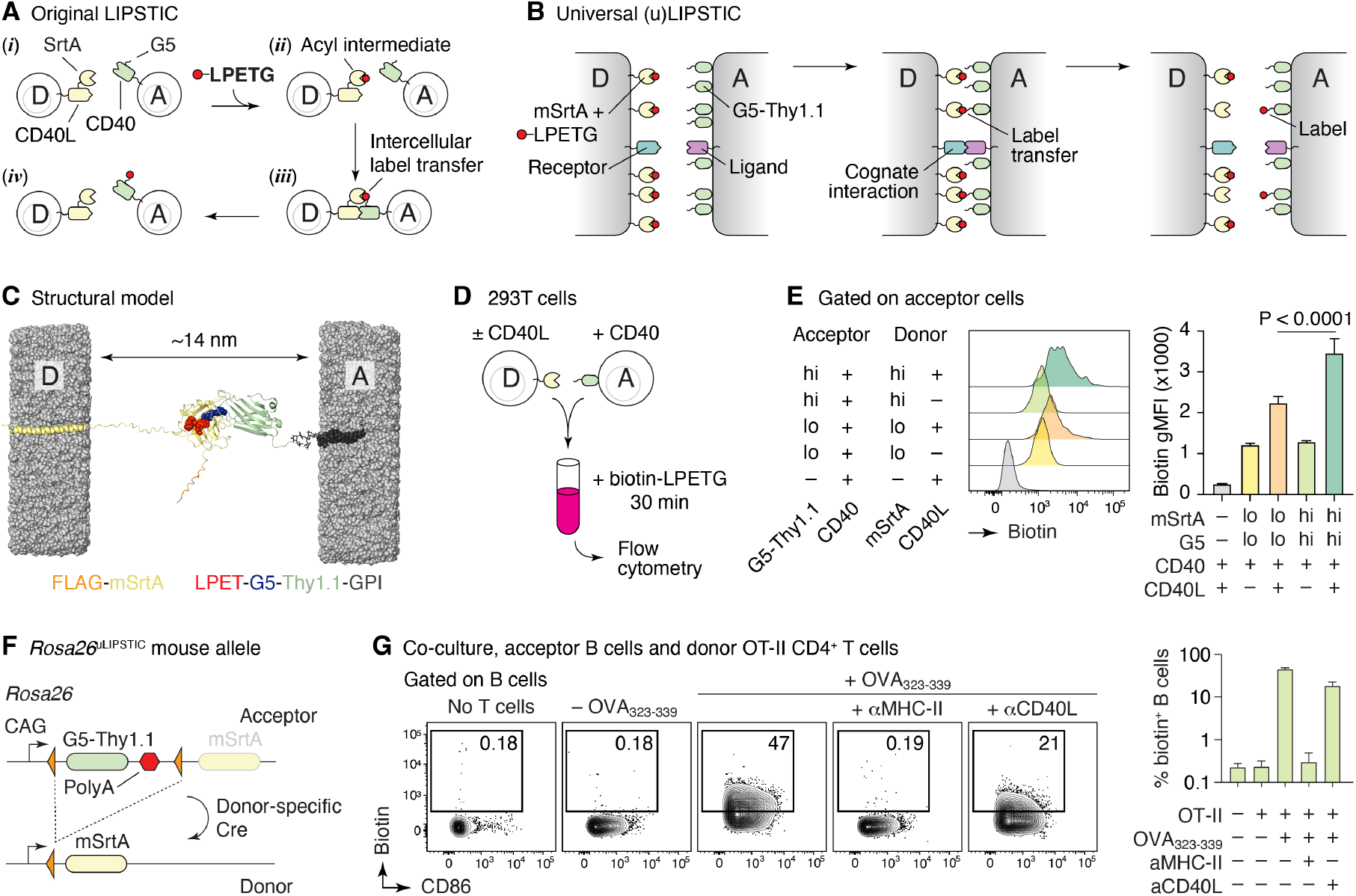
The uLIPSTIC system. **(A, B)** Schematic comparison of the original^24^ and universal LIPSTIC systems. In the original system (A), SrtA and G5 were brought into proximity by fusion to a receptor–ligand pair involved in a cell–cell interaction, allowing intercellular transfer of labeled substrate (LPETG) from donor cell “D” to acceptor cell “A.” In uLIPSTIC (B), SrtA and G5 (fused to the irrelevant protein Thy1.1) are anchored non-specifically to the cell membrane at high density; the enzymatic reaction is allowed to proceed when apposing membranes come within a short distance (< 14 nm) of each other, which can be driven by interactions between any receptor–ligand pair of the appropriate dimensions. **(C)** Computational model depicting the inter-membrane span of fully extended mSrtA upon transfer of the LPETG substrate onto G5-Thy1.1. **(D,E)** Populations of 293T cells co-transfected with high or low levels of either mSrtA or G5-Thy1.1 were co-incubated in the presence of biotin-LPETG for 30 min and analyzed by flow cytometry. **(F)** *Rosa26*^uLIPSTIC^ allele. Using the Ai9 high-expression backbone^32^, a LoxP-flanked G5-Thy1.1 is followed by mSrtA. Cre-recombinase switches cells from “acceptor” (G5-Thy1.1^+^) to “donor” (mSrtA^+^) modes. **(G)** *Rosa26*^uLIPSTIC/+^.CD4-Cre OT-II T cells were co-cultured with *Rosa26*^uLIPSTIC/+^ B cells in the presence or absence of OVA_323-339_ peptide and blocking antibodies to CD40L and MHC-II. Flow cytometry plots show biotin-LPETG transfer from T to B cells. Results from 2 independent experiments are summarized in the graph on the right.

### uLIPSTIC labeling of cell–cell interactions in vivo

To test the activity of uLIPSTIC *in vivo*, we used the antigen-specific interaction between DCs and CD4^+^ T cells. Since this interaction involves the CD40L–CD40 axis, we could benchmark the performance of uLIPSTIC against the original LIPSTIC system^24^. To this end, we used a classic *in vivo* T cell priming model^9, 33^, where G5-Thy1.1^+^ DCs loaded with OVA_323-339_ are injected into the footpads of mice followed by adoptive transfer of mSrtA^+^ OT-II T cells. Lymphatic migration of DCs to the draining popliteal lymph node (pLN) allows DC–T cell interactions to take place at this site (**Fig. 2A**). Footpad injection of biotin-LPETG substrate 24 h after T cell transfer led to detectable labeling of on average 6.5% of transferred DCs (**Fig. 2B**). Comparable numbers were obtained when using the original CD40L–CD40 LIPSTIC system^24^ (**Fig. 2C**). Treatment with anti-MHC-II prior to substrate injection blocked labeling in both settings (whereas treatment with anti-CD40L blocked transfer only by the original LIPSTIC), indicating that expression of the uLIPSTIC components alone is insufficient to artificially drive interactions between neighboring cells also *in vivo*. Thus, uLIPSTIC labeling is equivalent to receptor–ligand-specific LIPSTIC for recording the binding patterns of CD4^+^ T cells and DCs during *in vivo* priming. Of note, transferring increasing numbers of mSrtA^+^ donor T cells increased the degree to which interacting DCs were labeled (**Fig. S3A**). Thus, uLIPSTIC labeling efficiency varies with the availability of donor cells, likely as a consequence of multiple donor T cells being primed by the same acceptor DC.

**Figure 2.**
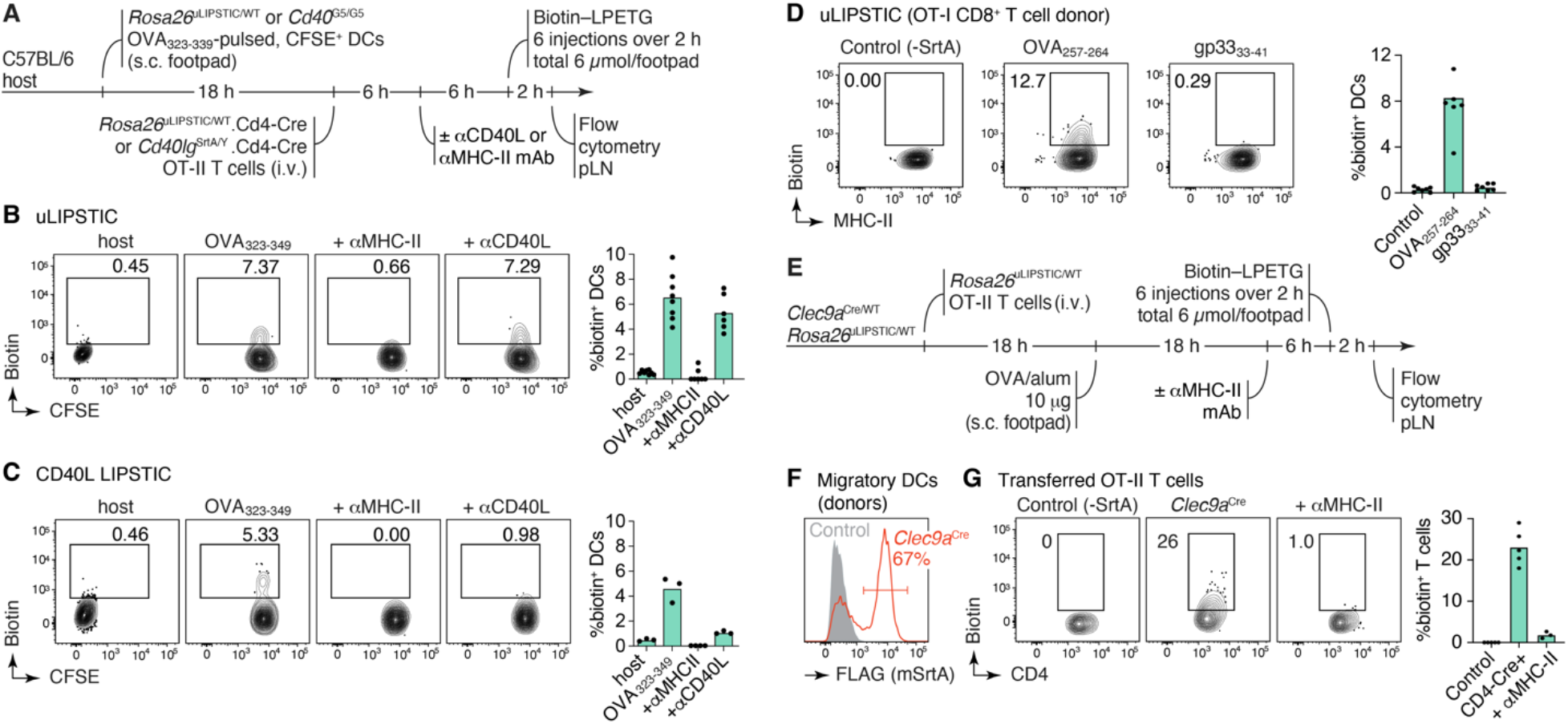
Bidirectional labeling of interactions between T cells and DCs in adoptive transfer models. **(A)** Experimental layout for the experiments in panels (B,C). **(B,C)** uLIPSTIC (B) and CD40L LIPSTIC (C) labeling of adoptively-transferred DCs in an *in vivo* priming model. Flow cytometry plots are gated on transferred (CFSE-labeled) DCs. Results from 2 independent experiments are summarized on the graphs to the right. **(D)** uLIPSTIC labeling of DCs by CD8^+^ T cells. Experimental setup as in (A), but DCs were pulsed either with cognate (OVA_257-264_) or control (LCMV gp33_33-41_) peptides. **(E-G)** Labeling of antigen-specific CD4^+^ T cells by *Clec9a*-expressing DCs. (E) Experimental layout. (F) efficiency of recombination of the uLIPSTIC allele in migratory (m)DCs by *Clec9a*^Cre^. (G) *Left*, labeling of adoptively transferred OT-II T cells upon immunization with OVA/alum. *Right*, summary of data from 2 independent experiments.

We next used uLIPSTIC to record interactions between T cells and DCs not accessible to the original LIPSTIC system, either because they do not involve the CD40L/CD40 interaction or because directionality is inversed. In an *in vivo* priming model, mSrtA^+^ OT-I CD8^+^ T cells were found to detectably label on average 14% of DCs pulsed with their cognate peptide (OVA_257-264_) but only background levels (0.6%) of DCs pulsed with a control peptide from lymphocytic choriomeningitis virus (**Fig. 2D**). We then inverted the uLIPSTIC reaction so that endogenous mSrtA^+^ DCs (in *Rosa26*^uLIPSTIC/+^.*Clec9a*^Cre/+^ mice^34^) were made to interact with adoptively transferred CD4^+^ (OT-II) T cells from Cre-negative *Rosa26*^uLIPSTIC/+^ donors upon immunization with OVA in alum adjuvant (**Fig. 2E**). This setup led to detectable labeling of roughly 20% of transferred T cells, which was again fully abrogated by prior injection of a blocking antibody to MHC-II (**Fig. 2F-G**). Thus, uLIPSTIC can label interactions between T cells and DCs bidirectionally.

To test the ability of uLIPSTIC to record immunological interactions other than priming of naïve T cells by DCs, we chose two settings. We first determined the identity of the cellular partners of regulatory T (Treg) cells in the steady state lymph node. To this end, we used the *Foxp3*^CreERT2^ allele^35^ to achieve tamoxifen-dependent recombination of *Rosa26*^uLIPSTIC^ specifically in Treg cells. We treated *Rosa26*^uLIPSTIC/+^.*Foxp3*^CreERT2/Y^ mice^35^ with tamoxifen to induce mSrtA expression in Treg cells, and then administered LIPSTIC substrate to footpads to achieve labeling in the draining pLNs (**Fig. 3A,B**). This revealed pronounced labeling of most DCs with migratory (MHC-II^hi^CD11c^int^) phenotype, whereas labeling of resident (MHC-II^int^CD11c^hi^) DCs was markedly lower (**Fig. 3B,C**). Importantly, labeling of Foxp3^−^ CD4^+^ T cells was negligible in this setting, confirming that simple colocalization of these cells with donor Treg cells within the same microenvironment is not sufficient to drive label transfer (**Fig. 3B, *left***). Expression of mSrtA^+^ in roughly equivalent numbers of Treg cells or total conventional CD4^+^ T cells (the latter achieved by low-dose tamoxifen administration to *Rosa26*^uLIPSTIC/+^.CD4-CreERT2 mice^36^) resulted in much less efficient labeling of migratory DCs by conventional T cells (**Fig. S3B,C)**. Thus, interaction with migratory DCs at steady state, although not a unique property of Treg cells, is more pronounced among this subset (**Fig. S3D**). Treg cell labeling of migratory DCs was decreased but not completely abrogated by administration of a blocking antibody to MHC-II, confirming that the strong interaction between Treg cells and migratory DCs is partly driven by the TCR-MHC-II axis but suggesting that other receptor-ligand pairs may also contribute to this process (**Fig. 3B,C**).

**Figure 3.**
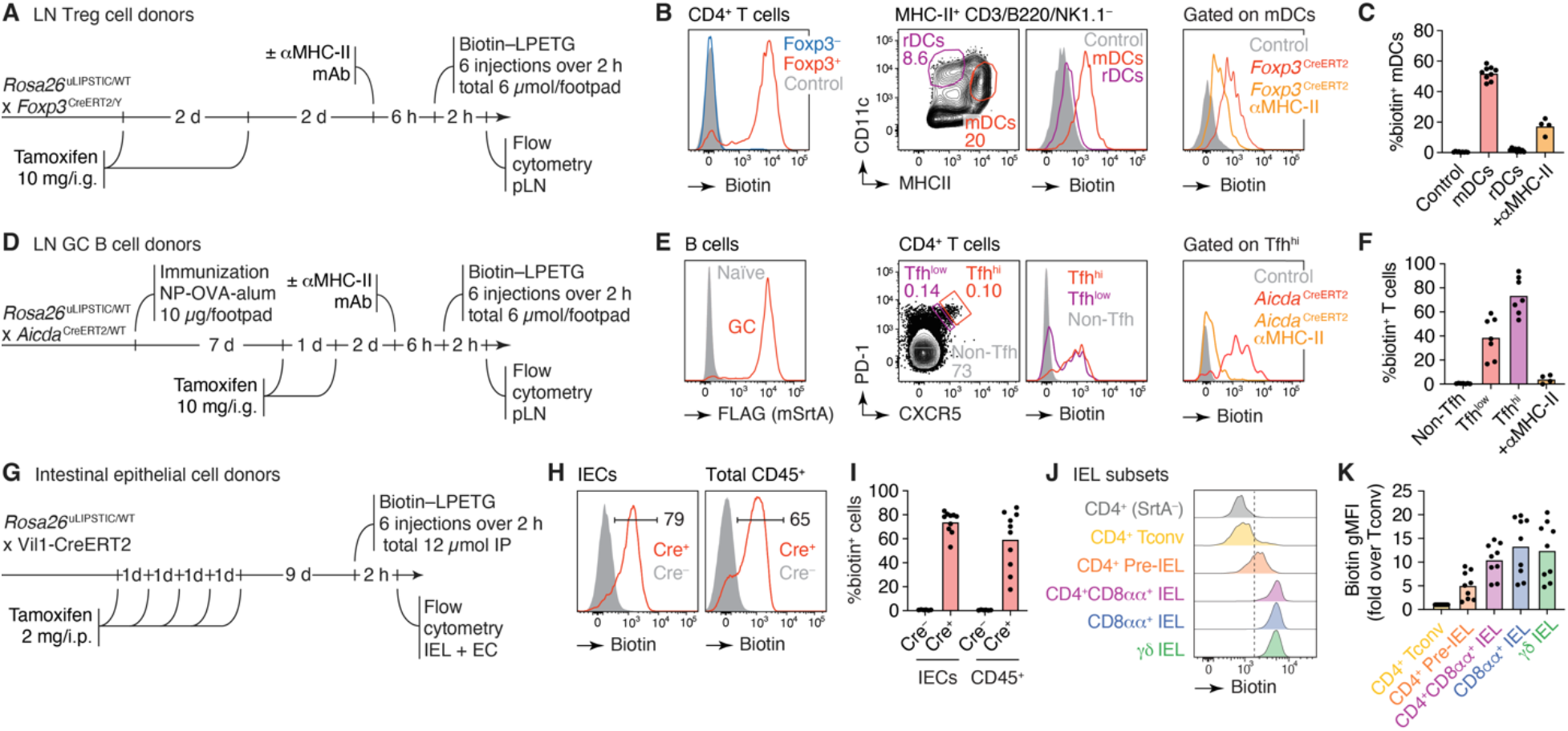
uLIPSTIC identifies cellular partners of Treg cells, Tfh cells, and IECs. **(A)** Experimental layout for panels (B,C). **(B)** *Left*, efficiency of recombination of the uLIPSTIC allele in Treg cells by *Foxp3*^CreERT2^. Biotin signal represents the acquisition of substrate by Treg cells (the biotin-LPET-SrtA acyl intermediate) and also shows the absence of transfer of substrate to *Foxp3*^−^ T cells. *Center*, labeling of migratory (m)DCs and resident (r)DCs by Treg cells at steady state. *Right*, labeling of mDCs upon injection of a blocking antibody to MHC-II. **(C)** Summary of data from 3 independent experiments. **(D)** Experimental layout for panels (E,F). **(E)** Labeling of Tfh cells by GC B cells. *Left*, efficiency of recombination of the uLIPSTIC allele in GC B cells by *Aicda*^CreERT2^ after 2 doses of tamoxifen, as in (B). *Center*, labeling of Tfh cells by GC B cells at 10 days after immunization with NP-OVA/alum. T cells are gated as high or low expressors of Tfh markers CXCR5 and PD-1 (Tfh^hi^ and Tfh^lo^, respectively). *Right*, labeling of Tfh^hi^ cells upon injection of a blocking antibody to MHC-II. **(F)** Summary of data from 2 independent experiments. **(G)** Experimental layout for panels (H-K). **(H)** Efficiency of conversion of IECs into uLIPSTIC donors and substrate capture in Vil1-CreERT2 mice (as in (B)) and transfer to total CD45^+^ intraepithelial leukocytes. **(I)** Summary of data from 4 mice in 2 independent experiments. **(J)** Differential labeling of selected IEL populations by IEC donors. The dashed line is placed for reference. **(K)** Summary of data as in (I). For all column plots, each symbol represents one mouse, bars represent the mean.

As a second test case, we used uLIPSTIC to determine the phenotype of the T cells that provide help to B cells in germinal centers (GCs), which are often difficult to identify unambiguously using the classic Tfh markers CXCR5 and PD-1^37^. To this end, we crossed *Rosa26*^uLIPSTIC^ to the *Aicda*^CreERT2^ allele, which allows for tamoxifen-dependent LoxP recombination in GC B cells^38^. We immunized *Rosa26*^uLIPSTIC/+^.*Aicda*^CreERT2/+^ mice^38^ in the footpads with the model antigen 4-hydroxi-3-nitro-phenylacetyl (NP)-OVA to generate GCs, then treated these mice with tamoxifen 7 and 8 days later to induce mSrtA expression in GC B cells (**Fig. 3D**). Footpad injection of biotin-LPETG 10 days post-immunization led to a high proportion of uLIPSTIC labeling among CXCR5^hi^PD-1^hi^ T follicular helper (Tfh) cells but not among CXCR5^−^PD-1^−^ non-Tfh cells in the pLN (**Fig. 3E,F**). Notably, only a fraction of T cells expressing lower levels of CXCR5 and PD-1 were labeled by GC B cells, indicating that relatively few of the cells within the “Tfh^low^” gate are truly engaged in helping GC B cells. Again, blocking of MHC-II led to total loss of Tfh cell labeling, confirming the specificity of the reaction (**Fig. 3E,F**).

Finally, to test the ability of uLIPSTIC to record interactions between immune cells and non-immune partners, we measured the transfer of substrate from intestinal epithelial cells (IECs) to the large population of intraepithelial T lymphocytes (IELs) that reside within this compartment^39^. We crossed *Rosa26*^uLIPSTIC^ mice to villin-1 (Vil1)-CreERT2^40^, a transgene that drives LoxP recombination in IECs upon administration of tamoxifen (**Fig. 3G**). Intraperitoneal administration of biotin-LPETG led to efficient loading of substrate onto IECs and transfer onto a large fraction (median 65%) of the CD45^+^ IEL population (**Fig. 3H,I** **and S4B**). Subsetting IELs into different populations showed that uLIPSTIC labeling followed a gradient corresponding to the stage of differentiation (or adaptation to the epithelial tissue) of these cells. Whereas labeling of “natural” TCRγ8^+^ and CD8αα^+^/TCRαβ^+^ IELs was uniformly high, uLIPSTIC signal in induced CD4^+^ IELs^41^ followed closely their developmental trajectory^5, 42^, from background levels in the CD4^+^CD8αα^−^CD103^−^ “conventional” subset to intermediate labeling in CD4^+^CD8αα^−^CD103^+^ pre-IELs and levels comparable to those of natural IELs in fully epithelium-adapted CD4^+^CD8αα^+^CD103^+^ population (**Fig. 3J,K**). Therefore, uLIPSTIC is capable of recording interactions between epithelial and immune cells in the small intestine. Our data show that close physical interaction with IECs is a common feature of most IELs, which is acquired in a stepwise fashion as CD4^+^ T lymphocytes differentiate into IELs and take up residence in the epithelium.

We conclude that uLIPSTIC can be used to label a wide variety of immune cell interactions *in vivo*, both in adoptive transfer and in fully endogenous models. In the latter, uLIPSTIC revealed the interaction preferences of steady-state LN Treg cells, identified populations of Tfh cells capable of providing help to B cells in the GC, and showed stepwise acquisition by intraepithelial CD4^+^ T cells of the ability to physically interact with IECs.

### Using uLIPSTIC for interaction-based transcriptomics

A key feature of uLIPSTIC is its ability to identify the full cellular interactome of a given cell type in an unbiased manner. Reading out this interactome is best achieved by single-cell RNA sequencing (scRNA-seq), which is also unbiased in its ability to identify labeled cell populations. Because LIPSTIC labeling has a wide dynamic range^24^, coupling it to scRNA-seq also has the potential to identify genes and transcriptional programs quantitatively associated with the degree of interaction between two cell types, which can in principle reveal the molecular pathways that drive a given interaction (**Fig. 4A**). To explore these possibilities, we profiled the immune interactome of IECs in the small intestine. *Rosa26*^uLIPSTIC/+^.Vil1-CreERT2 mice were labeled as in **Fig. 3G** and the intraepithelial immune cell (CD45^+^) fraction was stained with fluorescent antibodies for cell sorting plus a DNA-barcoded anti-biotin antibody to detect the uLIPSTIC signal. We sorted CD45^+^ immune cells in a semi-unbiased manner, using FACS gating to enrich for rarer leukocyte populations (**Fig. S4C**). These cells were then profiled by droplet-based scRNA-seq using the 10X Genomics platform. Immune cell populations were identified by comparing well characterized marker genes and publicly available gene signatures across the identified cell clusters, coupled with unbiased differential expression analysis and TCR sequence reconstruction (**Fig. 4B**, **Figs. S5-7, and Supplementary Spreadsheets 1,2**).

uLIPSTIC revealed broad variation in the extent to which different immune populations interact with IECs, which aligned with the data obtained by flow cytometry. Whereas there was a high degree of labeling among natural IEL subsets (TCRγ8 and TCRαβ^+^CD8αα^+^ lymphocytes), labeling was low or negligible among B cell populations (including naïve-like, activated, and plasma cell subsets), and intermediate in plasmacytoid DCs (**Fig. 4B**). uLIPSTIC also labeled two less clearly defined leukocyte clusters that interacted strongly with IECs. These included a small population of likely myeloid cells and a larger cluster marked by high expression of genes such as *Atxn1* and *Btbd11* (**Fig. 4C** **and Figs. S5A,G and S6A,B**). As seen by flow cytometry (**Fig. 3J,K**), CD4^+^ T cells again showed a gradient in their ability to interact with IECs, which became more apparent when these cells were clustered into subpopulations (**Fig. 4D**, **Fig. S7, and Supplementary Spreadsheet 3**). Acquisition of the biotin label largely followed a developmental trajectory that began with a highly polyclonal naïve-like population with low uLIPSTIC signal and followed through a pre-IEL intermediate into a fully differentiated, oligoclonal CD4-IEL state^5, 42^ (**Fig. 4D-F**) labeled to a similar extent as natural IELs (**Fig. 4C**).

**Figure 4.**
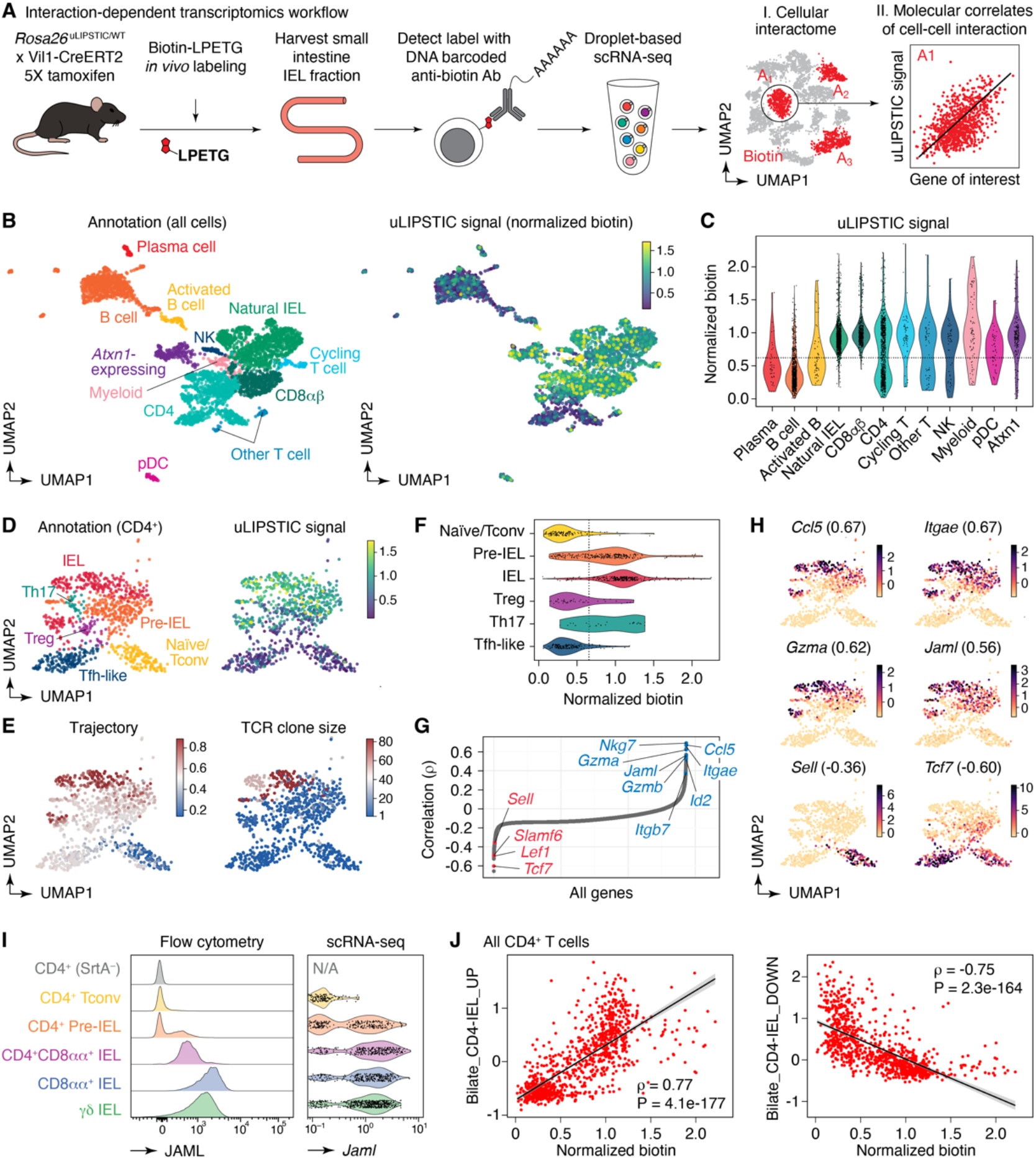
Using uLIPSTIC for interaction-based transcriptomics. **(A)** Experimental workflow. uLIPSTIC labeling is read out using DNA-barcoded antibodies and droplet-based scRNA-seq. Data are first queried for identification of acceptor (A_n_) populations, then uLIPSTIC signal within each population is correlated with expression of individual genes to search for mechanisms that drive donor-acceptor interaction. **(B)** UMAP plots of the CD45^+^ intraepithelial immune cell fraction from a uLIPSTIC reaction as in Fig. 3G. Data are pooled from three mice. *Left*, major cell populations (see **Fig. S5, S6**). *Right*, intensity of normalized uLIPSTIC signal (biotin). **(C)** Distribution of normalized uLIPSTIC signal among CD45^+^ cell populations. **(D)** UMAP plots of CD4^+^ T cells from (B). *Left*, major cell subpopulations (see **Fig. S7**). *Right*, intensity of normalized uLIPSTIC signal (biotin). **(E)** *Left*, inferred trajectory and *right*, αβTCR diversity (plotted as clone size) among CD4^+^ T cells. **(F)** Distribution of normalized uLIPSTIC (biotin) signal among CD4^+^ T cell subpopulations. **(G)** Correlation (Spearman’s *ρ*) between normalized uLIPSTIC signal and normalized gene expression, calculated for each gene over all CD4^+^ T cells, shown in order of increasing correlation. Selected genes are highlighted. Correlation for all selected genes was significant (FDR < 1e-23). **(H)** Normalized expression of selected genes. Correlation with normalized uLIPSTIC shown in parentheses. **(I)** Representative samples showing *in vivo* staining of JAML in IELs and scRNA-seq expression of *Jaml* in the equivalent populations. In the latter, CD8αα^+^ and γδ IEL were separated from within the “Natural IEL” cluster by the presence of rearranged αβ TCRs or expression of the *Trdc* gene. Fluorescent antibody was injected 6 h prior to IEL harvesting and analysis. **(J)** Correlation between normalized uLIPSTIC signal among all CD4^+^ T cells and expression of gene signatures up and downregulated as epithelial T cells transition from Tconv (CD4^+^CD103^−^CD8αα^−^) to CD4-IEL (CD4^+^CD103^+^CD8αα^+^) phenotypes (signatures based on data from Bilate et al.^42^). Trend line and error are for linear regression with 95% confidence interval.

To test the ability of uLIPSTIC transcriptomics to detect genes and pathways associated with the IEL–IEC interaction, we calculated the correlation between the uLIPSTIC signal within the CD4^+^ T cell population and the expression of all detected genes in our dataset (**Fig. 4G,H**). We found multiple significant correlations with key markers of IEL differentiation, including negative correlations with naïve T cell markers such as *Sell* (encoding for L-selectin) and *Tcf7* and positive correlations with the CD4-IEL associated genes such as *Ccl5*, *Gzma*, *Itgae*, *Itgb7*, and *Jaml* **(Fig. 4G,H, Fig. S7E,F, Supplementary Spreadsheet 4)**^42^. The last three genes are of particular interest, given that CD103 (the α_E_β_7_ integrin, encoded by *Itgae* and *Itgb7*) and JAML (junction adhesion molecule-like, encoded by *Jaml*) are interacting partners of E-cadherin and of the coxsackie and adenovirus receptor (CAR), respectively, both of which are expressed in the tight junctions of the intestinal epithelium^43–46^. In agreement with gene expression data, flow cytometry confirmed the correlation between biotin acquisition and expression of CD103 (**Fig. S8A**), and *in vivo* staining with an anti-JAML antibody confirmed the stepwise acquisition of this adhesion molecule during CD4-IEL development and its expression in multiple IEL subsets (**Fig. 4I**). Search for correlations among “canonical” (M2.CP) pathways in the MSigDB database^47^ revealed a significant positive correlation between biotin acquisition by CD4^+^ T cells and expression of genes in the Biocarta cytotoxic T lymphocyte (CTL) pathway, among others (**Fig. S8B**). Because many of the genes upregulated by CD4-IELs as they develop are related to the CTL program, we performed a targeted correlation analysis between biotin acquisition and the set of genes that change in expression as conventional T cells develop into CD4^+^ IELs^42^ (**Supplementary Spreadsheet 5**). This analysis revealed strong positive and negative correlations (|Spearman’s *ρ*| > 0.75) with genes up and downregulated, respectively, as CD4^+^ T cells differentiate towards the CD4-IEL fate (**Fig. 4J, Fig. S8C**). We conclude that uLIPSTIC can be used to achieve quantitative interaction-based transcriptomics, enabling us not only to define the “cellular interactomes” of populations of interest, but also to discern specific genes and gene signatures associated with acquisition of the ability to form specific cell-cell interactions.

## Discussion

This study describes a generalization of the LIPSTIC method applicable to cells of any type that interact physically with each other. Unlike the original version of our method^24^, uLIPSTIC does not require cognate interaction between a pre-specified receptor-ligand pair for label transfer, allowing one to probe the full cellular interactome of a population of interest in an unbiased manner. In the absence of such a requirement, the specificity of labeling is ensured by two properties of our system. First, the interacting components span a maximum intermembrane distance of ∼14 nm, typical of many receptor–ligand interactions including that of TCR with MHC^31^. Second, the extremely low affinity (millimolar Km) between the SrtA-LPETG donor and the G_5_ acceptor renders the uLIPSTIC components unlikely to themselves initiate unwarranted interactions between neighboring cells. Such specificity is supported by two separate lines of evidence. First, labeling is abrogated when antibodies are used to block known molecular drivers of a cellular interaction, demonstrating that efficient labeling takes place only when cells are allowed to interact functionally, such as through antigen presentation. Second, not all cells that are physically juxtaposed interact to the same extent, as exemplified by the low degree of labeling, if any, of conventional T cell or resident DC acceptors by Treg cell donors. uLIPSTIC thus differs from and complements methods such as synNotch variants, which, although they can be used to drive transcription of downstream reporter genes^26^, are based on molecular partners that bind each other with much higher (nanomolar) affinity, and thus are in theory themselves capable of driving cellular interactions^48^; as well as methods based on label transfer by extracellular diffusion^25^, which can better be conceived of as approaches to mark close cellular neighborhoods rather than functional interactions. uLIPSTIC also has advantages over cell doublet-based methods^49^, in that uLIPSTIC labeling is continuous rather than binary, can record cell–cell interactions that are not strong enough to survive cell extraction from tissue and flow cytometry processing, and does not require computational deconvolution of single-cell transcriptional profiles from doublets, although our system has the relative disadvantage of requiring genetic engineering of its components.

A central feature of uLIPSTIC is that it can be coupled directly to droplet-based scRNA-seq to achieve quantitative interaction-based transcriptomics. This property can be used in both an “atlas” mode, where the objective is to identify which populations of acceptor cells interact with a given donor lineage, and in “mechanistic” mode, where correlations between uLIPSTIC signal intensity and expression of individual genes or gene signatures allow us to establish the molecular basis of an interaction of interest. Using the second approach, we find that the ability of CD4^+^ T cells to interact physically with IECs in the small intestine is acquired developmentally, as these cells adapt to the intestinal tissue environment and acquire the phenotypic and transcriptional features of CD4-IELs^5, 42, 50^.

In conclusion, uLIPSTIC provides an unbiased platform for measurement of known cell-cell interactions as well as discovery of new ones. When coupled to scRNA-seq, uLIPSTIC interaction-based transcriptomics has the ability to quantify correlations between the intensity of cell-cell interactions and gene expression, allowing insight into the biology of the interaction itself. We expect this tool will be broadly useful for studying cellular interactions in immunology and beyond.

## Supporting information

Differentially expressed genes per Leiden cluster

Differentially expressed genes per annotated cluster

Differentially expressed genes between all cells within the CD4 cell type with all cells not in the cell type

Correlations between individual genes and acquisition of uLIPSTIC label among CD4+ T cells

Signatures of CD4-IEL and Tconv cells from Bilate et al, 2020

Sequences of DNA constructs used in this study

## Acknowledgements

We would like to thank the Rockefeller University Transgenics and Gene Targeting facilities for generating the uLIPSTIC mouse strain, the Comparative Biosciences Center for mouse housing, Tiago B.R. Castro for bioinformatics assistance, Kristie Gordon for cell sorting, and all Rockefeller University staff for their continuous support. We thank Caetano Reis e Sousa (Francis Crick Institute, UK), Tomohiro Kurosaki (U. Osaka, Japan), Takaharu Okada (RIKEN-Yokohama, Japan), and Claude-Agnès Reynaud and Jean-Claude Weill (Université Paris-Descartes, France) for mice; and Telmo Catarino (Instituto de Investigação e Inovação em Saúde, Portugal) and Henrique Veiga-Fernandes (Champalimaud Institute, Portugal) for their preliminary contributions to the development of the uLIPSTIC system.

## Author contributions

S. N.-H. carried out all experimental work with assistance from M.C.C.C., A.C., J.T.J., J.B., and S.M.P. S.W. and Y.P. designed and carried out all computational analyses. K.F. generated the structural model of the SrtA–G5-Thy1.1 interaction. G.P. conceived the uLIPSTIC system along with G.D.V. Y.P., A.M.B., E.F., D.M. and G.D.V. supervised the work. S.N.-H., S.W., Y.P., and G.D.V. wrote the manuscript, with input from all authors.

## Data availability

Final scRNA-seq datasets will be deposited to GEO upon publication. Processed scRNA-seq data and python code and Jupyter notebooks used for data analysis are available at https://github.com/pritykinlab/ulipstic-analysis.

## Methods

### Plasmids

All constructs were cloned into the pMP71 vector^51^, which was modified to express a fluorescent reporter (eGFP or tdTomato) followed by the porcine teschovirus-1 self-cleavable 2A peptide^52^ and the protein of interest. The SrtA sequence, including an N-terminal Flag-tag, was attached by a single Gly-Gly-Gly-Gly-Ser linker^53^ to the human PDGFRB transmembrane domain to form mSrtA. The pentaglycine (G5) acceptor sequence was fused at the N terminus of the mouse Thy1.1 protein, downstream of the signal peptide. Sequences of all constructs are included in **Supplementary Spreadsheet 5.**

### Mice

CD45.2 (C57BL6/J), CD45.1 (B6.SJL *Ptprc*^a^), CD4-Cre^54^, CD4-CreERT2^54^, and *Foxp3*^eGFP-CreERT2 35^ mice were purchased from the Jackson Laboratories (strain numbers 000664, 002014, 022071, 022356, and 016961, respectively). *Clec9a*^Cre^ mice^55^ were a kind gift from C. Reis e Sousa (Francis Crick Institute, UK), S1pr2-CreERT2 BAC-transgenic mice^56^ were generated and kindly provided by T. Kurosaki and T. Okada (Osaka University and RIKEN-Yokohama), and *Aicda^CreERT2^* mice^38^ were a kind gift from Claude-Agnès Reynaud and Jean-Claude Weill (Université Paris-Descartes). OT-II TCR transgenic (Y chromosome)^57^ mice were bred and maintained in our laboratory. The *Rosa26*^uLIPSTIC^ mouse strain was generated by the Rockefeller University Gene Targeting and Transgenics facilities, as described below. All genetically modified strains are bred and maintained under specific pathogen-free conditions at the Rockefeller University’s Comparative Biosciences Center in accordance with institutional guidelines and ethical regulations. All protocols were approved by the Rockefeller University Institutional Animal Care and Use Committee. Male and female 5–12-week-old mice were used in all experiments.

### Generation of the Rosa26^uLIPSTIC^ allele

*Rosa26^uLIPSTIC^* mice were generated by gene targeting in C57BL/6 embryonic stem cells (ESCs). The *Rosa26^uLIPSTIC^* targeting vector is a modification of the Ai9 *Rosa26* conditional expression vector^32^ (Addgene plasmid #22799). G_5_-Thy1.1 cDNA preceded by a mouse CD40 signal peptide was inserted into a NruI enzyme site in Ai9 immediately downstream of the first loxP site, whereas FLAG-mSrtA cDNA was introduced in place of the tdTomato gene using FseI enzyme sites. Expression of the cassette in ESCs was screened by standard Southern blotting analysis after EcoRI digestion and using a ^32^P probe targeting a sequence near the promoter region, shortly upstream of the left homology arm. Positive ESCs (7.3 kb band) were karyotyped, injected into blastocysts and chimeric founders were backcrossed to the C57BL6 background for at least six generations. The full sequence of the uLIPSTIC targeting vector and the Southern blot probe is reported in **Supplementary Spreadsheet 5.** uLIPSTIC mice were deposited at Jackson Labs under strain number 038221.

### Isolation of splenic dendritic cells (DCs), CD4^+^ T cells, and B cells

To isolate DCs, spleens were collected, incubated for 30 min at 37 °C in HBSS (Gibco) supplemented with CaCl2, MgCl2, and collagenase D at 400 U ml^-1^ (Roche). After digestion, tissue was forced 5 times through a 21-gauge needle and filtered through a 70 µm strainer into a 15 ml falcon tube with PBS supplemented with 0.5% BSA and 2 mM EDTA (PBE). Red-blood cells were lysed with ACK buffer (Gibco), and the resulting cell suspensions were filtered through a 70-μm mesh into PBE. DCs were obtained by magnetic cell separation (MACS) using anti-CD11c beads (Miltenyi Biotec), as per the manufacturer’s instructions. To isolate CD4^+^ T cells, CD8^+^ T cells, and B cells, spleens were processed as above, without collagenase digestion. CD4^+^ T cells and CD8^+^ T cells were isolated by negative selection using a cocktail of biotinylated antibodies targeting Ter119, CD11c, CD11b, CD25, B220, NK1.1, and either CD8 (for CD4^+^ isolation) or CD4 (for CD8^+^ isolation), followed by anti-biotin beads (Miltenyi Biotec), as per the manufacturer’s instructions. B cells were obtained by negative selection using anti-CD43 beads (Miltenyi Biotec), as per the manufacturer’s instructions.

### Adoptive cell transfers

For DC transfer experiments, splenic DCs were isolated as described above from mice subcutaneously injected with 1 × 10^6^ B16 melanoma cells that constitutively secrete FMS-like tyrosine kinase 3 ligand (Flt3L)^58^ 10 days prior to harvest. Cells were resuspended at 10^7^ cells/ml and incubated with 10 μM OVA^323–339^, LCMV-GP^61–80^, OVA^257-264^, or LCMV^276-286^ peptides (Anaspec) in RPMI + 10% FBS, for 30 min at 37 °C. For cell labelling, CFSE or CTV (ThermoFisher Scientific) was added to a final concentration of 2 μM during the last 5 or 20 minutes of incubation, respectively. Cells were washed three times in RPMI + 10% FBS and resuspended at 2 × 10^7^ cells/ml in PBS supplemented with 0.4 μg ml^−1^ LPS (Sigma-Aldrich). DCs were injected (5 × 10^5^ cells in 25 μl) subcutaneously into the hind footpads. For CD4^+^ T cell and CD8^+^ T cell transfer experiments, T cells isolated as described above were injected intravenously in 100 μl PBS per mouse.

### Immunizations

Mice were immunized by subcutaneous injection into the hind footpad of 10 μg OVA or 10 μg NP-OVA (Biosearch Technologies) adsorbed in alum (Imject Alum, ThermoFisher Scientific) at 2:1 antigen:alum (v:v) ratio in 25 μl volume.

### Antibody treatments

For CD40L and MHC Class II blocking experiments *in vivo*, mice were injected intravenously with 200 μg of CD40L-blocking antibody (clone MR-1, BioXCell) or subcutaneously with 150 μg of MHC-II (I-A/I-E) blocking antibody (clone M5/114, BioXCell), four hours prior to the first injection of substrate.

### Tamoxifen treatment

For induction of SrtA expression in regulatory T cells and conventional T cells, *Foxp3*^eGFP-CreERT2/Y^*.Rosa26*^uLIPSTIC/WT^ mice and CD4-CreERT2*.Rosa26*^uLIPSTIC/WT^ mice, respectively, were given two intragastric doses of 10 mg of tamoxifen (Sigma-Aldrich) dissolved in corn oil (Sigma-Aldrich) at 4 and 2 days prior to the end point (day 0). For SrtA expression in germinal center B cells, S1pr2-CreERT2.*Rosa26*^uLIPSTIC/WT^ mice and *Aicda*^CreERT2/+^.*Rosa26*^uLIPSTIC/WT^ mice, two doses of 10 mg of tamoxifen were administered at 3 and 2 days prior to the end point. SrtA expression in gut epithelial cells was induced by daily intraperitoneal injections of tamoxifen (2 mg per injection) for 5 consecutive days, starting 14 days before the end point.

### In vivo substrate administration

Biotin-aminohexanoic acid-LPETGS, C-terminus amide at 95% purity (biotin-LPTEG) was purchased from LifeTein (custom synthesis) and stock solutions were prepared in PBS at 20 mM. For *in vivo* LIPSTIC and uLIPSTIC labeling experiments in popliteal lymph nodes (pLNs), biotin–LPETG was injected subcutaneously into the hind footpad (20 μl of 2.5 mM solution in PBS) six times 20 min apart, and pLNs were collected 40 min after the last injection, as described^59^. Mice were briefly anaesthetized with isoflurane at each injection. For *in vivo* labeling of gut IELs, biotin-LPETG substrate was injected intraperitoneally (i.p.) (100 μl of 20 mM solution in PBS) six times 20 min apart. Small intestines were collected 40 min after the last injection.

### Isolation of lymphocytes from pLNs

pLNs were collected into microfuge tubes with 500 μl HBSS (Gibco) supplemented with CaCl_2_, MgCl_2_, and collagenase D at 400 U ml^-1^ (Roche). pLNs were cut into small pieces and incubated for 30 min at 37 °C . After digestion, tissue was forced 5 times through a 21-gauge needle and filtered through a 70 µm strainer into a 15 ml falcon tube with PBE.

### Isolation of intraepithelial leukocytes

Intraepithelial leukocytes were isolated as previously described ^60^. Briefly, small intestines were harvested and washed in PBS. Peyer’s patches were surgically removed prior to incubation with 1 mM dithiothreitol followed by 30 mM EDTA. Intraepithelial cells were recovered from the supernatant of dithiothreitol and EDTA washes and mononuclear cells were isolated by 40% and 80% gradient Percoll centrifugation.

### Flow Cytometry and cell sorting

Single-cell suspensions were washed with PBE, incubated with 1 μg ml^−1^ anti-CD16/32 (2.4G2, BioXCell) for 5 min at room temperature and then stained for cell surface markers at 4 °C for 20 min in PBS using the reagents listed in **Supplementary Table 1**. Cells were washed with PBE and stained with Zombie fixable viability dye (BioLegend) or fixable Aqua dead cell stain kit (Invitrogen) at room temperature for 15 min, then washed with PBE and filtered through a 40 μm strainer for acquisition. For *in vivo* JAML staining of IELs, mice were injected i.p. with 100 μg of anti-JAML AF646 antibody 12 or 6 h prior to the end point. For single-cell transcriptomic analysis, stained cells were further incubated with DNA-barcoded anti-biotin and sample hashtag (anti-MHC-I) antibodies (BioLegend) for 20 minutes in PBE, washed three times with PBE, and bulk-sorted. For substrate detection *in vivo*, an anti-biotin–PE antibody (Miltenyi Biotec) was exclusively used, as described previously^59^. Samples were acquired on FACS Symphony or Fortessa analyzers or sorted on FACSAriaII or FACSAriaIII cell sorters (BD Biosciences). Data were analyzed using FlowJo v.10.6.2 software.

### uLIPSTIC labeling in vitro

HEK293T cells (ATCC) were transfected by calcium phosphate transfection with the indicated expression vectors at high (1 μg μl^-1^) and low (0.1 μg μl^-1^) concentrations of Thy1.1-G5 and mSrtA constructs. Forty hours after transfection, cells were detached TrypLE Express cell dissociation solution (ThermoFisher Scientific), washed and resuspended at 10^6^ cell per ml in PBS. Donor cell populations transfected with CD40L and/or mSrtA constructs and acceptor cell populations transfected with CD40 and/or Thy1.1-G5 were mixed at a 1:1 ratio (10^5^ cells of each population) in a 1.5-ml conical tube, to which biotin–LPETG was added to a final concentration of 100 μM. Cells were incubated at room temperature for 30 min and washed three times with PBE to remove excess biotin–LPETG before FACS staining.

### uLIPSTIC labeling ex vivo

B cells from *Rosa26*^uLIPSTIC/WT^ mice and CD4^+^ T cells from OT-II CD4-Cre.*Rosa26*^uLIPSTIC/WT^ mice were isolated from mouse spleens as described above. Isolated T cells were activated with CD3/CD28 dynabeads (ThermoFisher Scientific) for 24 h and then co-cultured with isolated B cells (2 × 10^5^ cells per well, 1:1 ratio) in the presence or absence of OVA^323–339^ peptide in RPMI, 10% FBS supplemented with 0.1% 2 mercaptoethanol (Gibco) in U-bottom 96-well plates for 20 h. Blocking antibodies were added at the beginning of the co-culture at a final concentration of 150 μg ml^−1^. To label interactions ex-vivo, biotin-LPETG substrate was added 30 min before harvest at a final concentration of 100 μM.

### Library preparation for single cell-RNA sequencing of IELs

In addition to fluorescent antibodies, cells were co-stained prior to sorting with hashtag oligonucleotide (HTO)-labeled antibodies to CD45 and MHC-I for sample separation (two hashtags per sample) and HTO-anti-biotin for detection of the uLIPSTIC signal. Sorted cells were collected into a microfuge tube with 300 μl PBS supplemented with 0.4% BSA. After the sort, tubes were topped with PBS 0.4% BSA, centrifuged and the buffer was carefully reduced by removing the volume with a pipette to a final volume of 40 μl. Cells were counted for viability and immediately submitted to library preparation. The scRNA-seq library was prepared using the 10X Single Cell Chromium system, according to the manufacturer’s instructions, at the Genomics Core of Rockefeller University and was sequenced on an Illumina NovaSeq SP flowcell to a minimum sequencing depth of 30,000 reads per cell using read lengths of 26 bp read 1, 8 bp i7 index, 98 bp read 2.

### Single-cell RNA sequencing analysis

Gene expression unique molecular identifier (UMI) counts, along with sample and biotin (uLIPSTIC) HTO counts, were generated with CellRanger v6.0.1 “count” using “Feature Barcode” counts and otherwise default parameters, with mm10 reference. TCR data were preprocessed with CellRanger “vdj” with default parameters. Applying default cellranger filtering, this resulted in a filtered gene expression UMI count matrix including 4,607 cells and 32,285 genes.

The scanpy package v1.9.1 was used for all analysis of the gene expression data^61^. Cell barcodes with unresolved sample HTOs, a low or extremely high number of expressed genes, a large fraction of expressed mitochondrial genes, or likely doublets were removed. Genes expressed in a low number of cells were removed. This resulted in a filtered gene expression matrix of 3,677 cells and 14,332 genes with a matching biotin HTO count in each cell representing uLIPSTIC signal.

Gene counts were normalized using Pearson residual normalization with theta = 1. Principal component analysis (PCA) was run with default parameters, and then k nearest neighbor (kNN) graph was constructed using 40 PCs, k = 30 and otherwise default parameters. Then Leiden clustering was performed with a resolution of 1, resulting in 26 clusters.

uLIPSTIC normalized values were obtained for each cell by dividing the uLIPSTIC HTO counts by the number of sample-encoding HTO read counts in a cell. The 5th percentile of these normalized values was added as a pseudocount, and then log10 applied. These values were then shifted by the minimum log-scaled value, so the scale starts at 0.

Known marker genes as well as TCR data were used to annotate the Leiden clusters. The scirpy package v0.10.1 was used for the TCR data preprocessing and analysis^62^. Cluster 10 was split into two subclusters that contained cycling T and B cells. Annotations were confirmed by scoring PanglaoDB immune cell marker gene sets^63^ using the score_genes() function in scanpy and by exploring significantly differentially expressed genes in each cluster as compared with all cells outside the cluster, obtained using a custom script. For differential expression analysis, log2 fold change (log2FC) of expression was calculated as the ratio of pseudobulk raw UMI counts summed over cells within and outside the cluster (then normalized by total amount of UMI counts inside and outside the cluster), p-values were calculated using Mann-Whitney U test applied to Pearson residual normalized expression values in single cells within and outside the cluster, and Bonferroni correction for multiple hypothesis testing applied to all genes with abs(log2FC) > 0.5.

The analysis then focused on the CD4 T cell subset. A new kNN graph was generated for this subset, again using k = 30 neighbors and 40 PCs, and Leiden clustering performed with resolution = 1.3. Clusters were annotated using known marker genes and TCR clonality information. Trajectory analysis and subsequent cell pseudotime calculation were performed using Wishbone v0.5.2^64^ using default parameters as available in scanpy and using as the root the cell in the Naive/Tconv cluster with the highest value of *Sell* expression.

To identify candidate genes involved in cell-cell interactions, for every gene the Spearman correlation was calculated between the Pearson residual normalized value of expression of that gene and the uLIPSTIC signal across all cells in the CD4 T cell subset. Bonferroni correction was used for multiple hypothesis testing on all genes. This calculation was separately performed when removing Tfh-like and naïve/memory cells, or when restricting to cells from each individual mouse, with consistent results.

For violin plot of scRNA-seq expression of *Jaml* (**Fig. 4I**), Pearson residual normalized values were shifted so that the minimum value is zero, bottom 5^th^ percentile of all values (across cell groups) omitted, and then plotted on log scale. The T cell subpopulations for the plot were defined as follows. The subpopulations of CD4 T cells, “Naïve/Tconv”, “Pre-IEL”, “IEL” (**Fig. 4D-H, S7**), were used as CD4^+^ Tconv, CD4^+^ Pre-IEL and CD4^+^ CD8αα^+^ IEL, respectively. The “Natural IEL” cells (**Fig. 4B, S5, S6**) were separated into three groups: CD8αα^+^ IEL if TCR ab chain was detected (301 cells), otherwise γδ IEL if normalized expression of Trdc was above 0 (517 cells), and other (163 cells) which were not included in the plot.

MSigDB canonical pathways were scored using scanpy’s score_genes() function over all CD4+ T cells. Spearman correlation with normalized biotin values was calculated for all pathways, and p-values were adjusted using q-value approach^65^ for pathways with positive correlation values. Top 5 pathway scores are shown by correlation value, for those with q < 0.05 **(Fig. S8B)**. CD4^+^CD103^+^CD8αα^+^ and CD4^+^CD103^−^CD8αα^−^ gene signatures were generated from scRNA-seq (library 2) from Bilate et al^42^. tdTomato^−^CD4^+^CD8αα^+^ cells (Cluster 2) were compared to tdTomato^−^ “recent epithelial immigrants” (REI, Cluster 5) using the Wilcoxon Rank Sum test. P-values were adjusted using Bonferroni correction. All genes with adjusted p-values < 0.05 were included in the signature. Genes with positive fold-change (enriched in Cluster 2) were included in “Bilate_CD40-IEL_UP” and with negative fold-change (enriched in Cluster 5) were included in “Bilate_CD40-IEL_DOWN”. Signatures were scored on the uLIPSTIC scRNA-seq data using scanpy’s score_genes() function, and Spearman correlation with normalized biotin values for both gene signatures was calculated over all CD4+ T cells, or over all CD4+ T cells excluding Tconv and Tfh-like cells. Linear regression fit with 95% confidence interval overlayed over scatter plots was calculated using geom_smooth() in ggplot2 using default parameters.

### Modelling the SrtA-Thy1.1 complex on cells surfaces

First, structures of G5-Thy1.1 and FLAG-SrtA-PDGFRb were generated using Alphafold2^66^. Next, in the FLAG-SrtA-PDGFRB model, the domain constructing peptide binding domain was substituted with the substrate bound Sortase A structure (PDB: 1T2W). Additionally, the flexible linker connecting SrtA domain to PDGFRB transmembrane helix was rebuilt to an extended conformation using COOT^67^ to better estimate the maximum distance the protein is able to extend to. The Thy1.1 was aligned to SrtA using the substrate of 1T2W and 5G acceptor motif of G5-Thy1.1. Any resulting interprotein clashes were corrected using GalaxyWEB server^68^. To build the GPI anchor and the lipid bilayers we used CHARMM-GUI^69^. The anchor glycolipid was generated based on the human prion protein (PrP) GPI^70^. Next theFLAG-SrtA-PDGFRB:G5-Thy1.1 complex was modelled in the POPC/cholesterol lipid bilayer using CHARMM-GUI. The GPI anchor was placed in the second bilayer using ChimeraX^71^. Finally, both bilayers, with the protein complex and the GPI anchor were aligned in ChimeraX. The distance between two bilayers was measured in PyMOL^72^.

### Statistical analysis

Statistical tests were performed in GraphPad Prism 9.0 software. Comparisons between two treatment conditions were analyzed using unpaired, two-tailed Student’s t-test and multivariate data were analyzed by one-way ANOVA with Tukey’s post-hoc tests to further examine pairwise differences.

## Supplementary Figures

**Supplementary Figure 1.**
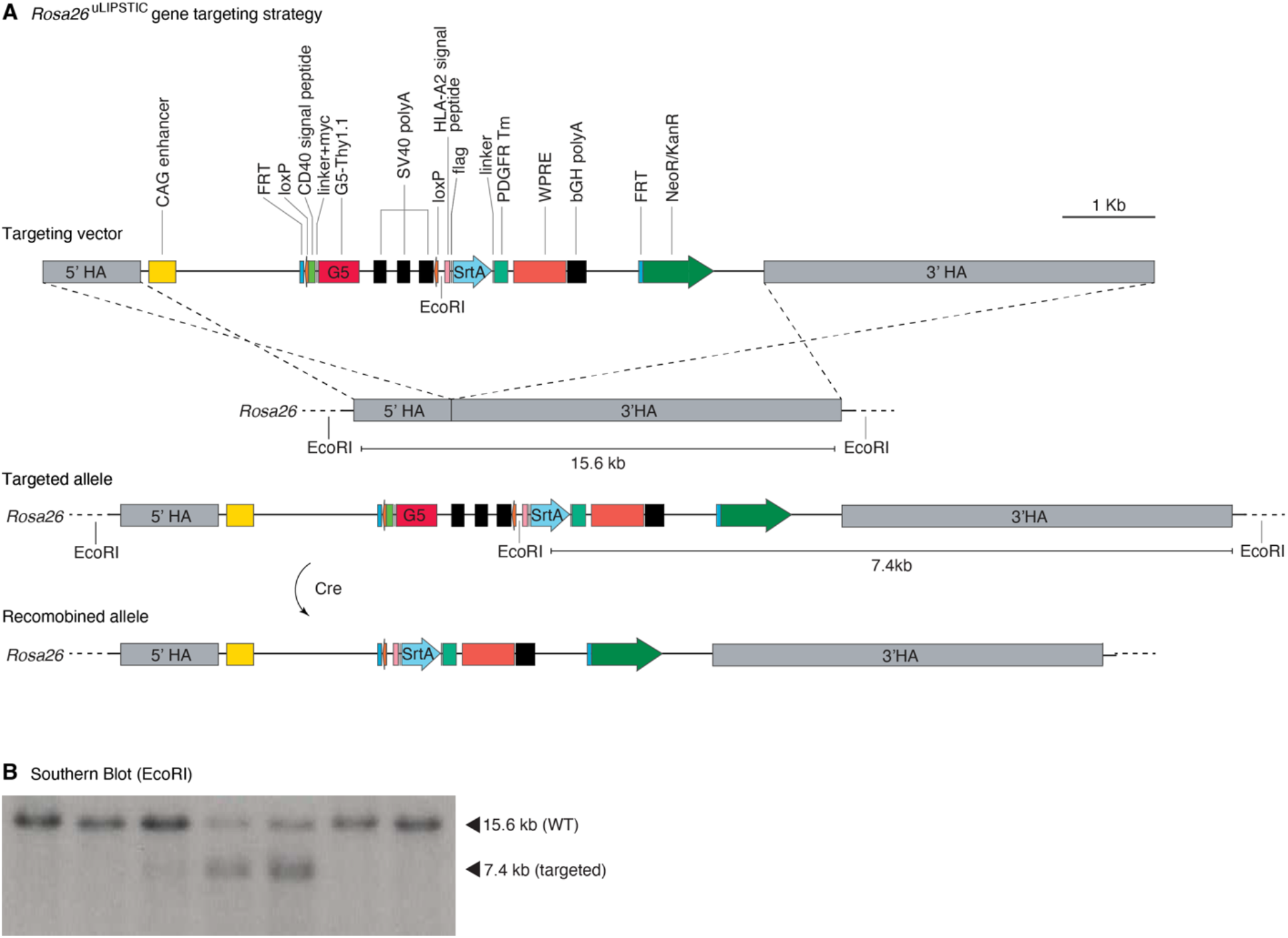
uLIPSTIC design *in vivo*. **(A)** The uLIPSTIC cassette carrying the lox-stop-lox G5Thy1.1 followed by mSrtA-PDGFRtm was cloned into the Ai9 *Rosa26* targeting plasmid. **(B)** Insertion of the uLIPSTIC cassette was assessed in embryonic stem (ES) cells by Southern blotting using a ^32^P-labeled probe (**Supplementary Spreadsheet 6**) annealing upstream of the left arm after EcoRI digestion. ESCs carrying the insertion exhibit an extra EcoRI restriction site, resulting in a 7.4 kb fragment upon enzymatic digestion. The blot shows 2 heterozygous integrations out of 7 ES cell clones screened.

**Supplementary Figure 2.**
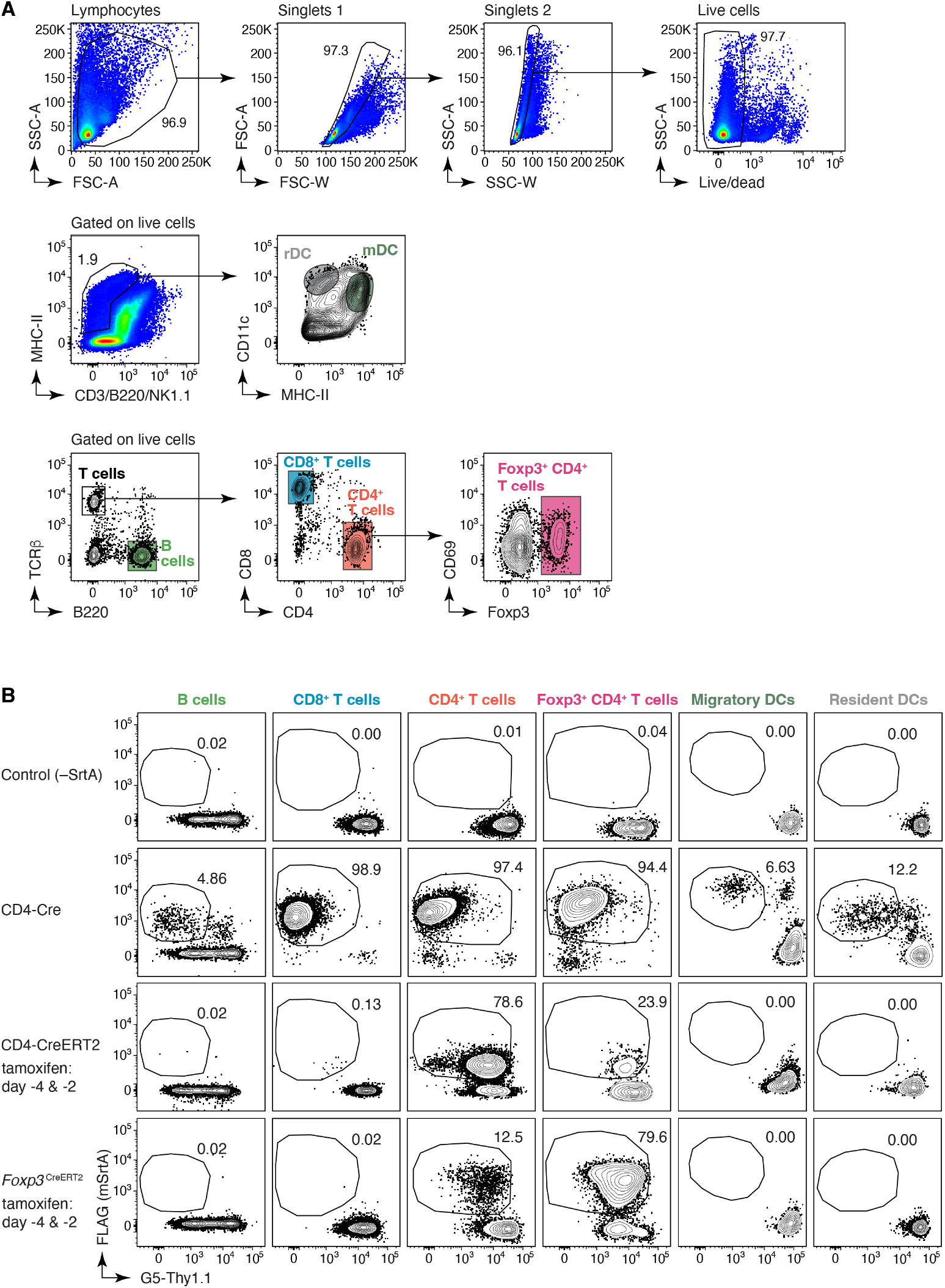
SrtA expression in donor cells is determined by the specificity of the cre driver. **(A)** Representative gating strategy for resident dendritic cells (rDCs; LIN^−^, MHC-II^int^, CD11c^hi^), migratory dendritic cells (mDCs; LIN^−^, MHC-II^hi^, CD11c^+^), CD4^+^ T cells, CD8^+^ T cells, (Treg) cells and B cells in lymph nodes. **(B)** SrtA expression (determined by FLAG detection) is induced by Cre recombination. Use of a constitutive Cre line (e.g., CD4-Cre) results in efficient but non-specific SrtA expression, generating T cells that can only be used in adoptive cell transfer experiments. The use of inducible Cre lines such as CD4-CreERT2 and *Foxp3*^CreERT2^ can often resolve specificity issues, enabling the implementation of uLIPSTIC in fully endogenous models.

**Supplementary Figure 3.**
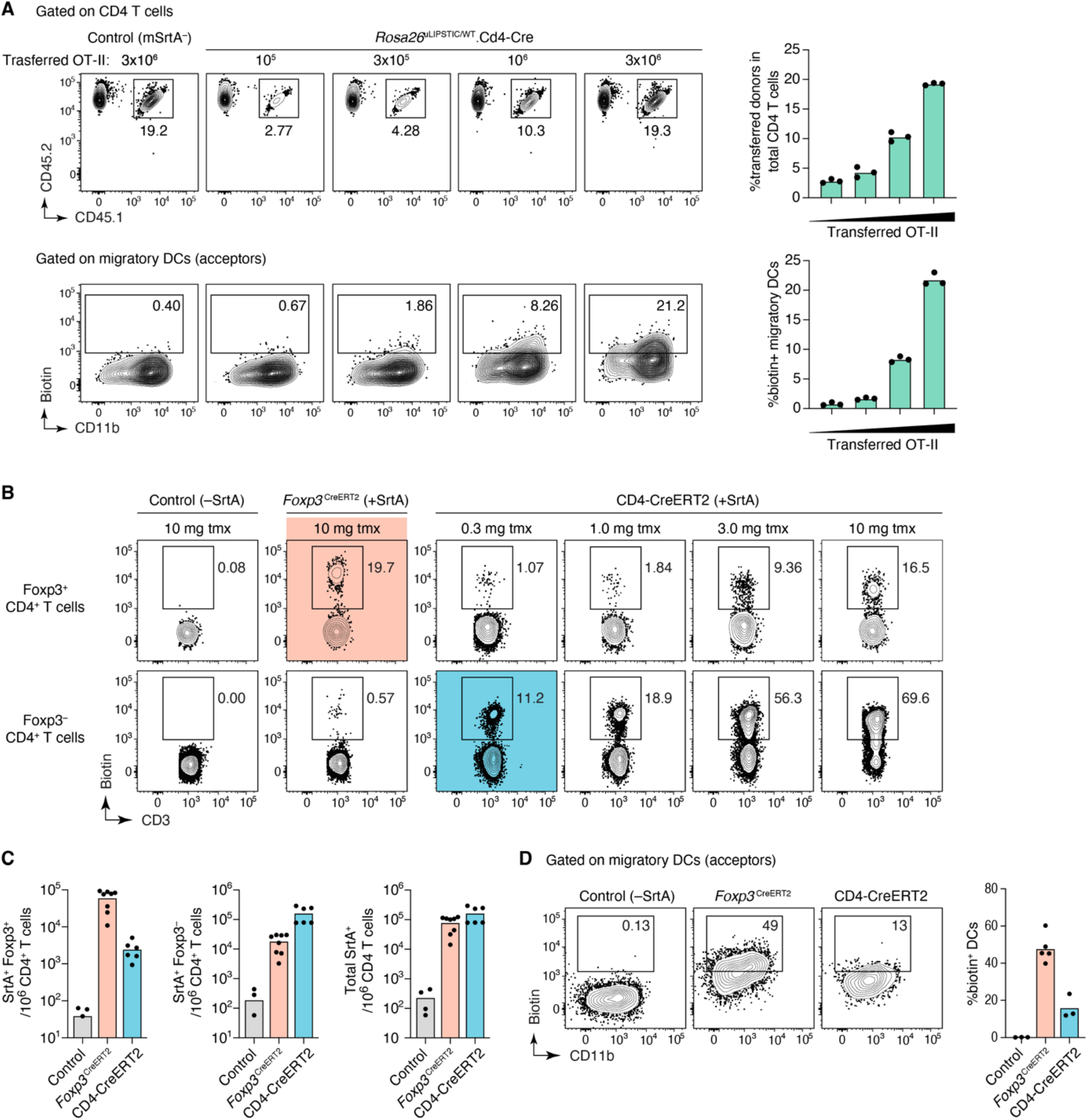
uLIPSTIC labeling of T cell–DC interactions. **(A)** mSrtA^+^ donor cell numbers determine the degree of uLIPSTIC labeling. Increasing numbers (10^5^, 3 – 10^5^, 10^6^, 3 – 10^6^) of *Rosa26*^uLIPSTIC/+^.CD4-Cre OT-II CD4+ T cells were adoptively transferred into recipient *Rosa26*^uLIPSTIC/ uLIPSTIC^ mice, followed by OVA/alum immunization 18 h post-transfer and LIPSTIC substrate injection one day later. The number of transferred cells (CD45.1/2) determined the proportion of donor cells in the CD4^+^ T cell compartment (*top*) and the corresponding percentage of labeled interacting cells in the mDC compartment (*bottom*). **(B-D)** Treg cells preferentially interact with mDCs (B-D). (B) To test if enhanced interaction with mDCs is a specific feature of Treg cells or a general feature of all CD4^+^ T cells, we titrated the dose of tamoxifen in *Rosa26*^uLIPSTIC/+^.CD4-CreERT2 mice to achieve a similar percentage of SrtA-expression among total CD4^+^ T cells as in *Rosa26*^uLIPSTIC/+^.*Foxp3*^CreERT2/Y^ mice. (C) At a dose of 0.3 mg of tamoxifen, *Rosa26*^uLIPSTIC/+^.CD4-CreERT2 mice showed SrtA expression in a small number of Treg cells (*left*), with most SrtA^+^ cells observed in CD4^+^ conventional T cells (*center*) and overall numbers of SrtA^+^ cells among total CD4^+^ T cells that were comparable with those of *Rosa26*^uLIPSTIC/+^.*Foxp3*^CreERT2/Y^ mice treated with 10 mg tamoxifen (*right*). (D) When numbers of Treg and CD4^+^ conventional donor cells are equalized, acceptor mDCs show stronger interaction with Treg cell partners.

**Supplementary Figure 4.**
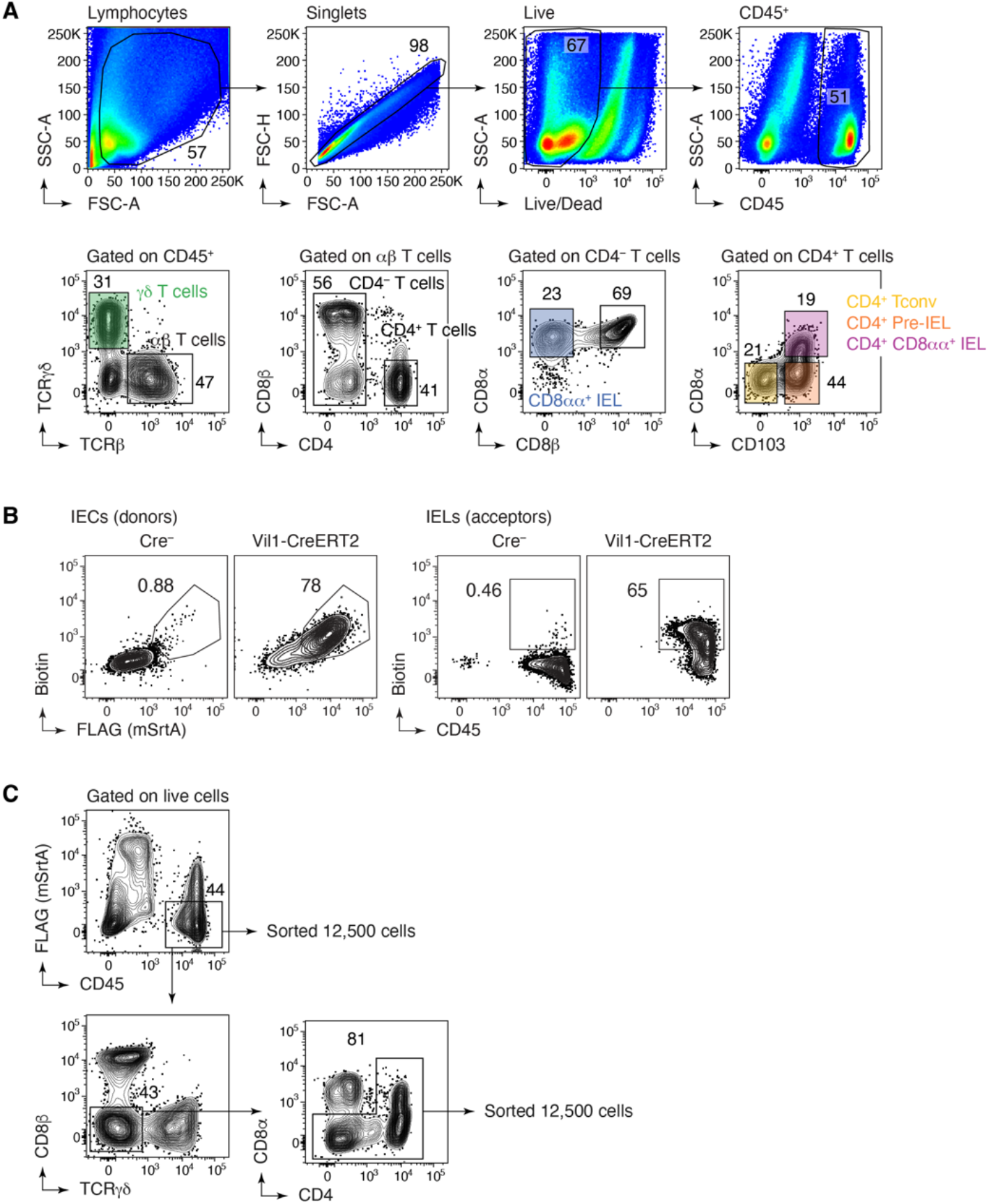
Flow cytometry strategy for intraepithelial immune cells. **(A)** Representative gating strategy for γ8 TCR and αβ TCR (Cd8αα^+^, CD8αβ^+^, and CD4^+^) IEL subsets. **(B)** *Left*, expression of SrtA (FLAG) and capture of LIPSTIC substrate by IEC donor cells and *right*, transfer of substrate onto CD45^+^ acceptor cells in SrtA-expressing and control mice. **(C)** Sorting strategy for the scRNA-seq experiment. Samples were enriched for rarer (e.g., B cell, CD4-IEL) populations by first sorting 12,500 total cells then an additional 12,500 cells depleted of the dominant γ8, CD8αα, and CD8αβ IEL populations. Three independent samples were sorted and stained with different hashtag oligos for downstream identification.

**Supplementary Figure 5.**
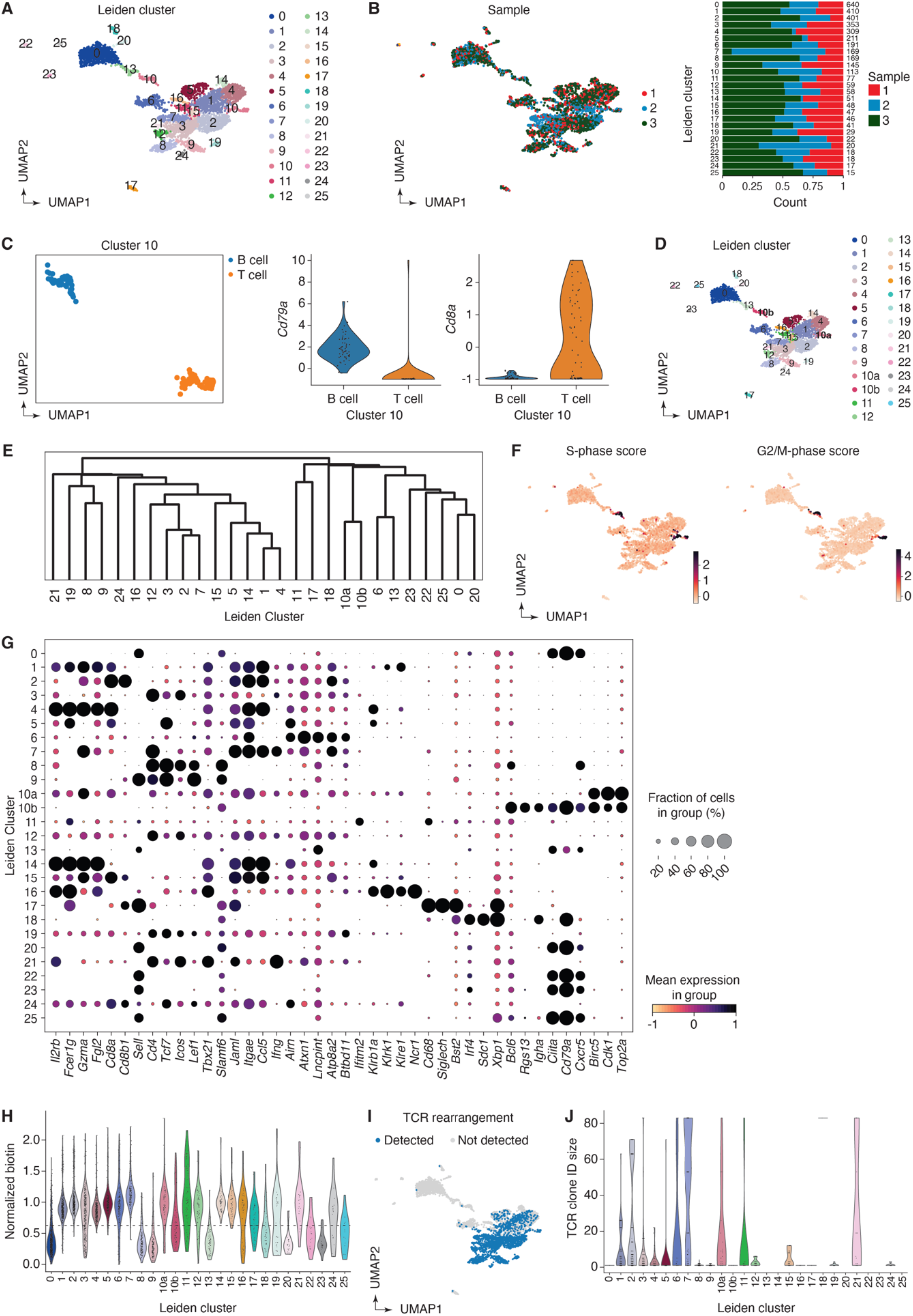
Clustering analysis of the immune interactome of IECs in the small intestine. **(A)** UMAP colored by Leiden clustering of the entire scRNA-seq/uLIPSTIC dataset (n=3,677 cells). **(B)** *Left*, UMAP colored by biological replicate. *Right*, bar plot indicating cluster composition by biological replicate, cluster size indicated at the right of each bar the right. **(C)** *Left*, UMAP for reanalysis and sub-clustering of Leiden cluster 10 into two clusters. *Right*, normalized expression of *Cd79a* and *Cd8a* for these two sub-clusters of cluster 10, determining their annotation as either B or T cells. **(D)** UMAP showing final clustering of the entire data. **(E)** Dendrogram representing transcriptional similarities among clusters. Differentially expressed genes were identified for each cluster (log2FC > 1, FDR < 0.05, see *Methods*), and normalized expression of all such genes (5,756 genes total), averaged per cluster, was used for the hierarchical clustering analysis that produced the dendrogram. **(F)** UMAP showing the S and G2M phase cell cycle gene list scores (obtained using the score_genes_cell_cycle() function with lists from the Seurat package^73^). **(G)** Dot plot of marker genes indicating their level of expression in each cluster. Dot size indicates the fraction of cells in the cluster with Pearson residual normalized expression greater than 0, dot color represents level of expression. **(H)** Violin plot showing levels of normalized uLIPSTIC signal for each Leiden cluster. **(I)** Left, UMAP showing presence of rearranged TCRα and β in each cell. Right, violin plot indicating the sizes of clones containing cells from each cluster.

**Supplementary Figure 6.**
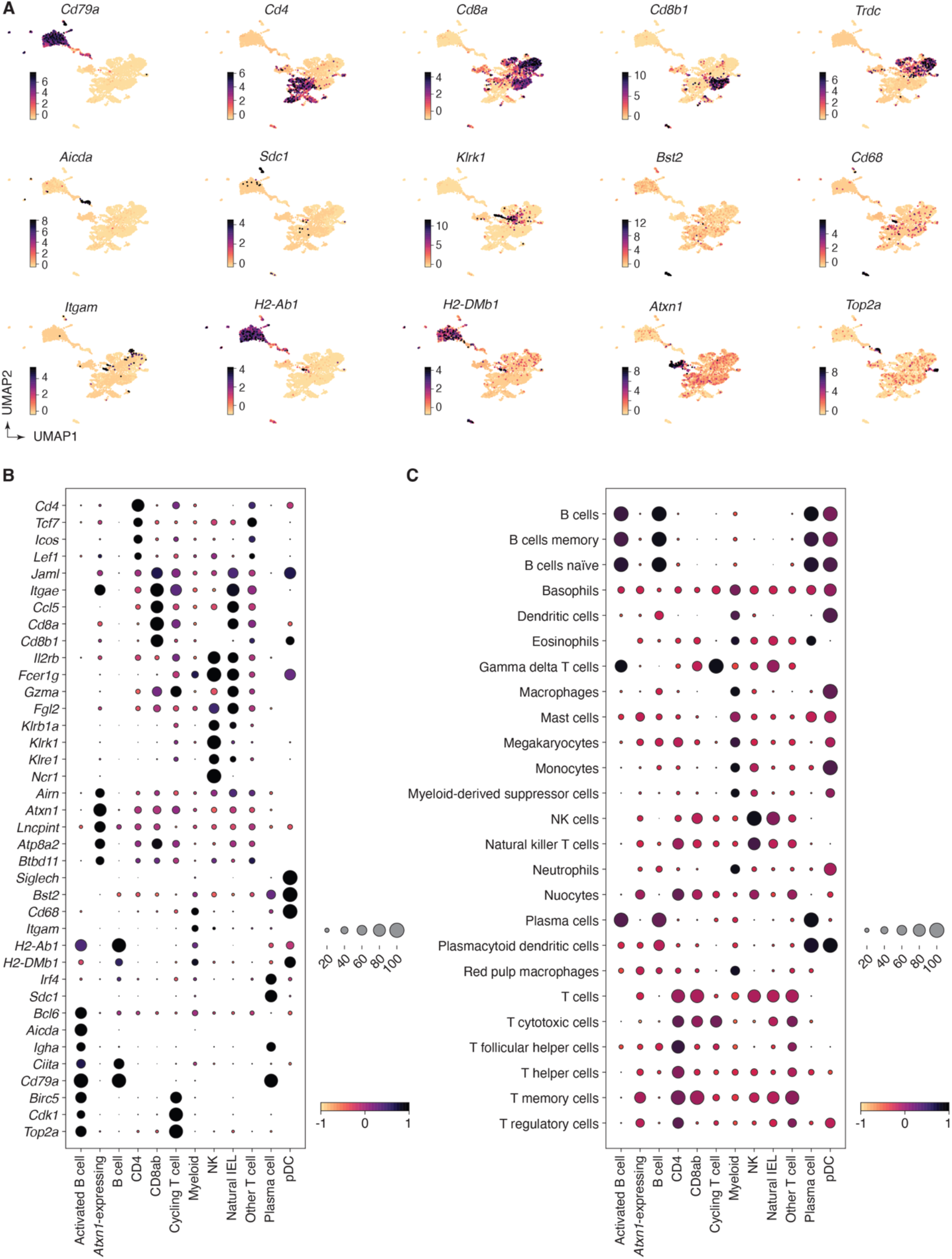
Expression of marker genes and gene signatures in the annotated scRNA-seq data. **(A)** UMAP plots showing normalized gene expression levels for selected marker genes. **(B)** Dot plot of marker genes indicating level of expression for each cell type annotation. **(C)** Dot plot of scores for gene signatures of immune cell types from PanglaoDB^63^. For both dot plots, dot size indicates the fraction of cells in the cluster with Pearson residual normalized expression greater than 0, dot color represents level of expression.

**Supplementary Figure 7.**
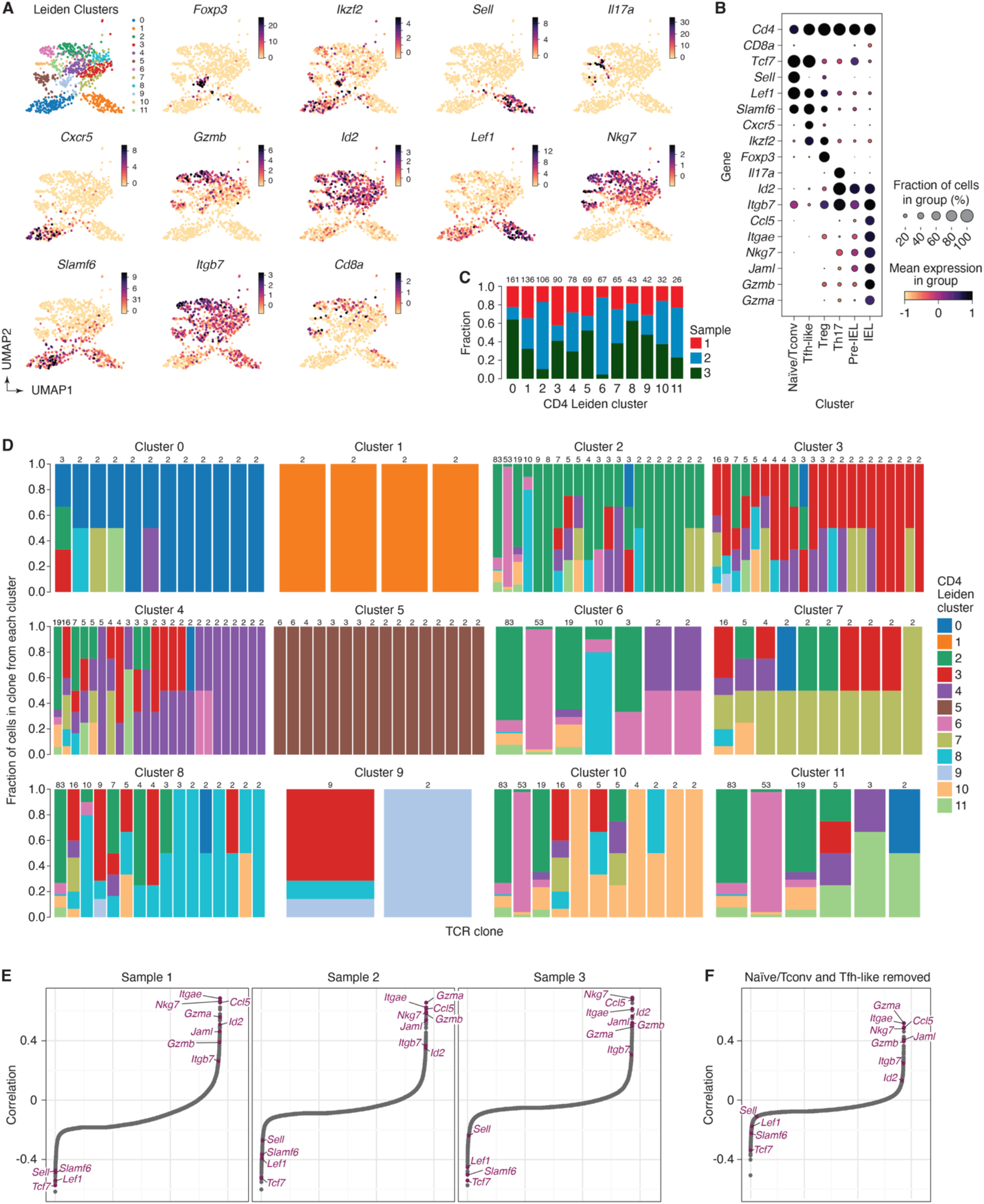
Analysis of combined scRNA-seq + uLIPSTIC data for CD4^+^ T cells. **(A)** UMAP for CD4^+^ T cells showing new Leiden sub-clusters and expression of selected marker genes in each cluster. **(B)** Dot plot of marker genes for each annotated subset of CD4^+^ T cells. Dot size indicates the fraction of cells in the cluster with Pearson residual normalized expression greater than 0, dot color represents level of expression. **(C)** Bar plot indicating CD4^+^ T cell sub-cluster composition by biological replicate, cluster size indicated at the right of each bar. **(D)** Bar plots showing the cluster composition of TCR clones. Each bar plot corresponds to a cluster (Leiden sub-clusters of CD4 T cells) and represents the composition for each clone (with at least two cells) such that at least one cell from that clone belongs to that cluster. Clone size is given on top of each bar. By definition, each clone selected for CD4 cluster 0 has at least one cell from CD4 cluster 0, each clone selected for cluster CD4 cluster 1 has at least one cell from CD4 cluster 1, etc. Each clone can therefore be represented in multiple bar plots. **(E)** Spearman correlation values, in increasing order, for uLIPSTIC signal and normalized expression of a gene, calculated separately for cells from each biological replicate. **(F)** Spearman correlation values, in increasing order, for uLIPSTIC signal and normalized expression of a gene, calculated when removing Tfh-like and naïve/conventional T cells (Leiden CD4 sub-clusters 0 and 1).

**Supplementary Figure 8.**
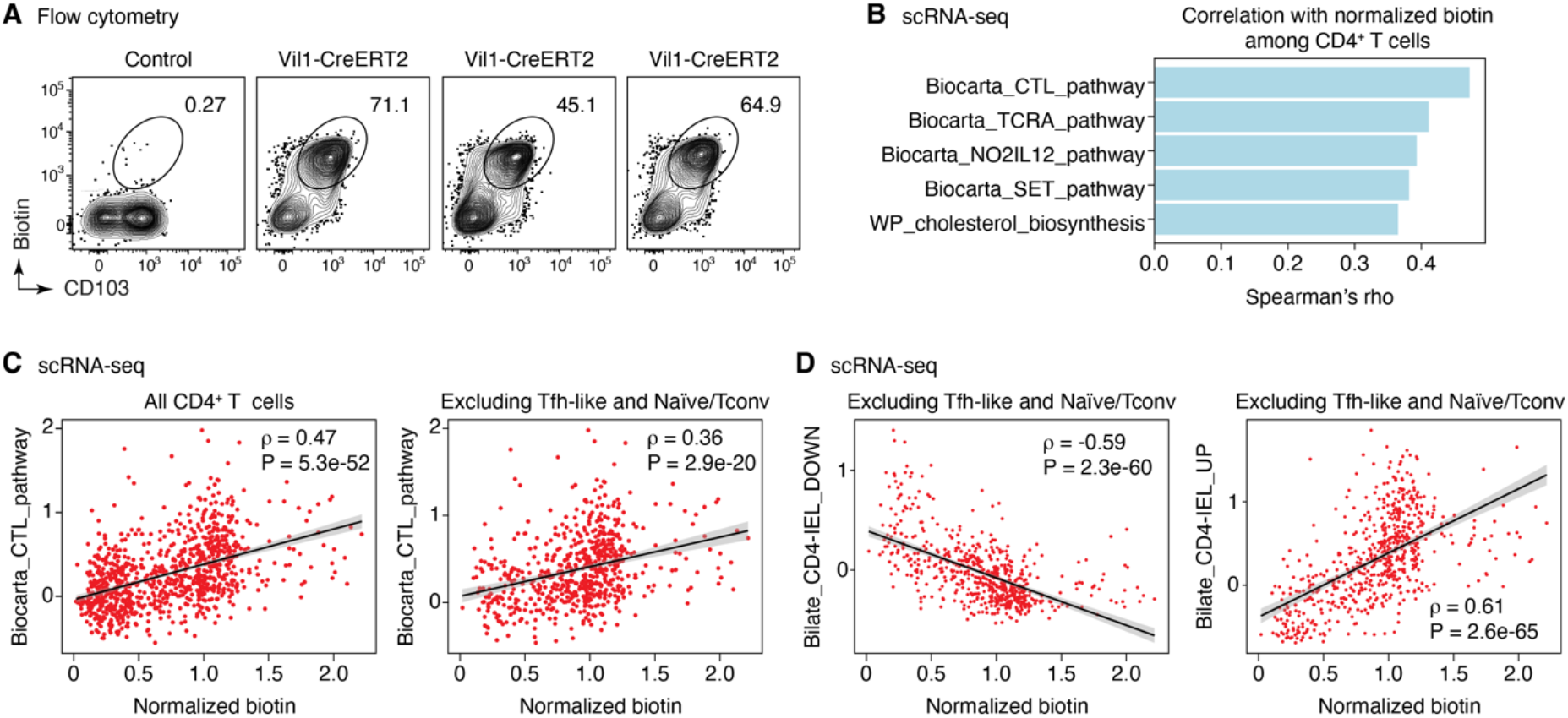
Correlation between acquisition of uLIPSTIC label and expression of CD103 and selected gene signatures by CD4-IELs. **(A)** Flow cytometry plots show uLIPSTIC signal and CD103 expression in one control *Rosa26*^uLIPSTIC/WT^ and three Vil1-Cre.*Rosa26*^uLIPSTIC/WT^ mice treated as in Fig. 3G. **(B)** Gene signatures from the MSigDB “canonical pathways” (M2.CP) database showing significant positive association with normalized biotin signal in scRNA-seq analysis over all CD4^+^ T cells. Plots show Spearman’s *ρ* value for each signature. **(C)** Correlation between acquisition of uLIPSTIC signal by CD4^+^ T cells (shown for all T cells and excluding Tfh-like and Naïve/Tconv clusters) and expression of the Biocarta CTL gene signature. Trend line and error are for linear regression with 95% confidence interval. **(D)** Correlation between acquisition of uLIPSTIC signal by CD4^+^ T cells (shown for T cells excluding Tfh-like and Naïve/Tconv clusters) and expression of gene signatures up and downregulated as epithelial T cells transition from Tconv (CD4^+^CD103^−^CD8αα^−^) to CD4-IEL (CD4^+^CD103^+^CD8αα^+^) phenotypes (signatures based on data from Bilate et al.^42^). Trend line and error are for linear regression with 95% confidence interval.

**Supplementary Table 1.** Antibodies used for Flow Cytometry, CITE-seq, and *in vivo* blocking.

| Antibody | Source | Identifier |
| --- | --- | --- |
| Rat monoclonal anti-MHC-II BV421 (clone M5/114.15.2) | BioLegend | 107632 |
| Rat monoclonal anti-MHC-II FITC (clone M5/114.15.2) | BioLegend | 107606 |
| Rat monoclonal anti-MHC-II AF700 (clone M5/114.15.2) | Invitrogen | 56-5321-82 |
| Rat monoclonal anti-MHC-II BUV395 (clone 2G9)v | BD Biosciences | 743876 |
| Hamster monoclonal anti-CD11c BUV496 (clone N418) | BD Biosciences | 750450 |
| Armenian hamster monoclonal anti-CD11c APC (clone N418) | BioLegend | 117310 |
| Rat monoclonal anti-CD11b BV711 (clone M1/70) | BioLegend | 101242 |
| Mouse monoclonal anti-XCR1 PerCP-Cy5.5 (clone ZET) | BioLegend | 148208 |
| Mouse monoclonal anti-biotin PE (clone Bio3-18E7) | Miltenyi Biotec | 130-090-756 |
| Mouse monoclonal anti-biotin APC (clone Bio3-18E7) | Miltenyi Biotec | 130-113-288 |
| Rat monoclonal anti-DYKDDDDK (FLAG) APC (clone L5) | BioLegend | 637308 |
| Rat monoclonal anti-DYKDDDDK (FLAG) BV421 (clone L5) | BioLegend | 637321 |
| Mouse monoclonal anti-CD90.1 (Thy-1.1) BV421 (clone OX-7) | BioLegend | 202529 |
| Mouse monoclonal anti-CD90.1 (Thy-1.1) PE-Cy7 (clone OX-7) | BioLegend | 202518 |
| Armenian hamster monoclonal anti-CD3ε BV785 (clone 145-2C11) | BioLegend | 100355 |
| Mouse monoclonal anti-CD19 BV605 (HIB19) | BioLegend | 302244 |
| Rat monoclonal anti-B220 BUV395 (clone RA3-6B2) | BD Biosciences | 563793 |
| Rat monoclonal anti-B220 FITC (clone RA3-6B2) | BioLegend | 103206 |
| Rat monoclonal anti-B220 BV785 (clone RA3-6B2) | BioLegend | 103245 |
| Rat monoclonal anti-NK1.1 BV785 (clone PK136) | BioLegend | 108749 |
| Rat monoclonal anti-F4/80 PE-Cy7 (clone BM8) | BioLegend | 123114 |
| Rat monoclonal anti-CD4 BV785 (RM4-5) | BioLegend | 100551 |
| Rat monoclonal anti-CD4 PE-Cy7 (GK1.5) | Invitrogen | 25-0041-81 |
| Rat monoclonal anti-CD4 BUV496 (GK1.5) | BD Biosciences | 612952 |
| Rat monoclonal anti-CD4 AF700 (RM4-5) | BD Biosciences | 557956 |
| Rat monoclonal anti-CD8α BUV805 (53-6.7) | BD Biosciences | 612898 |
| Rat monoclonal anti-CD45 FITC (30-F11) | BD Biosciences | 553080 |
| Rat monoclonal anti-CD45 AF700 (30-F11) | BioLegend | 103128 |
| Mouse monoclonal anti-CD45.1 PE-Cy7 (A20) | BioLegend | 110730 |
| Mouse monoclonal anti-CD45.2 FITC (104) | BioLegend | 109816 |
| Mouse monoclonal anti-CD45.2 PE-Cy7 (104) | BioLegend | 109830 |
| Rat monoclonal anti-CD86 BV605 (GL1) | BioLegend | 105037 |
| Rat monoclonal anti-CD86 AF488 (GL1) | BioLegend | 105018 |
| Rat monoclonal anti-CD40 BV421 (clone 3/23) | BD Biosciences | 562846 |
| Armenian hamster monoclonal anti-CD80 PerCP-Cy5.5 (clone 16-10A1) | BioLegend | 104722 |
| Armenian hamster monoclonal anti-TCRβ APC-eFluor780 (clone H57-597) | Invitrogen | 47-5961-82 |
| Hamster monoclonal anti-TCRβ BUV395 (clone H57-597) | BD Biosciences | 742485 |
| Armenian hamster monoclonal anti-CD103 BV785 (clone 2E7) | BioLegend | 121439 |
| Rat monoclonal anti-FOXP3 APC (clone FJK-16s) | Invitrogen | 17-5773-82 |
| Rat monoclonal anti-FOXP3 PerCP-Cy5.5 (clone FJK-16s) | Invitrogen | 45-5773-82 |
| Rat monoclonal anti-FOXP3 FITC (clone FJK-16s) | Invitrogen | 11-5773-82 |
| Armenian hamster monoclonal anti-JAML AF647 (clone 4E10) | BioLegend | 128506 |
| Armenian hamster monoclonal anti-TCRγ/δ PerCP-eFluor710 (clone GL3) | Invitrogen | 46-5711-82 |
| Rat monoclonal anti-CD326 (Ep-CAM) BV605 (clone G8.8) | BioLegend | 118227 |
| Rat monoclonal anti-CD16/CD32 (clone 2.4G2) | BioXCell | BP0307 |
| Rat monoclonal anti-CD8a Biotin (clone 53-6.7) | BioLegend | 100704 |
| Rat monoclonal anti-CD11b Biotin (clone M1/70) | BioLegend | 101204 |
| Mouse monoclonal anti-NK1.1 Biotin (clone PK136) | BioLegend | 108704 |
| Rat monoclonal anti-CD25 Biotin (clone 7D4) | BD Biosciences | 553070 |
| Armenian hamster monoclonal anti-CD11c Biotin (clone N418) | BioLegend | 117304 |
| Rat monoclonal anti-TER-119 Biotin (clone TER-119) | BD Biosciences | 553672 |
| Rat monoclonal anti-B220 Biotin (clone RA3-6B2) | BD Biosciences | 553086 |
| Rat monoclonal anti-CD4 Biotin (clone GK1.5) | BioLegend | 100404 |
| Armenian hamster monoclonal anti-CD40L (clone MR1) | BioXCell | BE0017-1 |
| Rat monoclonal anti-MHC-II (clone M5/114) | BioXCell | BE0108 |
| Armenian hamster monoclonal anti-JAML (clone 4E10) | BioLegend | 128502 |
| TotalSeq-C0301 Rat monoclonal anti-CD45 (clone 30-F11) and anti-MHC-I (clone M1/42) Hashtag 1 | BioLegend | 155861 |
| TotalSeq-C0302 Rat monoclonal anti-CD45 (clone 30-F11) and anti-MHC-I (clone M1/42) Hashtag 2 | BioLegend | 155863 |
| TotalSeq-C0303 Rat monoclonal anti-CD45 (clone 30-F11) and anti-MHC-I (clone M1/42) Hashtag 3 | BioLegend | 155865 |
| TotalSeq-C0304 Rat monoclonal anti-CD45 (clone 30-F11) and anti-MHC-I (clone M1/42) Hashtag 4 | BioLegend | 155867 |
| TotalSeq-C0305 Rat monoclonal anti-CD45 (clone 30-F11) and anti-MHC-I (clone M1/42) Hashtag 5 | BioLegend | 155869 |
| TotalSeq-C0306 Rat monoclonal anti-CD45 (clone 30-F11) and anti-MHC-I (clone M1/42) Hashtag 6 | BioLegend | 155871 |
| TotalSeq-C0307 Rat monoclonal anti-CD45 (clone 30-F11) and anti-MHC-I (clone M1/42) Hashtag 7 | BioLegend | 155873 |
| TotalSeq-C0308 Rat monoclonal anti-CD45 (clone 30-F11) and anti-MHC-I (clone M1/42) Hashtag 8 | BioLegend | 155875 |
| TotalSeq-C0436 Mouse monoclonal anti-biotin (clone 1D4-C5) | BioLegend | 409011 |

## References

1. Greenwald I, Rubin GM. Making a difference: the role of cell-cell interactions in establishing separate identities for equivalent cells. Cell. 1992;68(2):271–81. doi: 10.1016/0092-8674(92)90470-w. PubMed PMID: 1365402.

2. Sudhof TC, Malenka RC. Understanding synapses: past, present, and future. Neuron. 2008;60(3):469–76. doi: 10.1016/j.neuron.2008.10.011. PubMed PMID: 18995821; PMCID: PMC3243741.

3. Dustin ML. The immunological synapse. Cancer Immunol Res. 2014;2(11):1023–33. doi: 10.1158/2326-6066.CIR-14-0161. PubMed PMID: 25367977; PMCID: PMC4692051.

4. Victora GD, Nussenzweig MC. Germinal centers. Annu Rev Immunol. 2012;30:429-57. Epub 2012/01/10. doi: 10.1146/annurev-immunol-020711-075032. PubMed PMID: 22224772.

5. London M, Bilate AM, Castro TBR, Sujino T, Mucida D. Stepwise chromatin and transcriptional acquisition of an intraepithelial lymphocyte program. Nat Immunol. 2021;22(4):449–59. Epub 2021/03/10. doi: 10.1038/s41590-021-00883-8. PubMed PMID: 33686285; PMCID: PMC8251700.

6. Hoytema van Konijnenburg DP, Reis BS, Pedicord VA, Farache J, Victora GD, Mucida D. Intestinal Epithelial and Intraepithelial T Cell Crosstalk Mediates a Dynamic Response to Infection. Cell. 2017;171(4):783–94 e13. Epub 2017/09/26. doi: 10.1016/j.cell.2017.08.046. PubMed PMID: 28942917; PMCID: PMC5670000.

7. Niec RE, Rudensky AY, Fuchs E. Inflammatory adaptation in barrier tissues. Cell. 2021;184(13):3361–75. doi: 10.1016/j.cell.2021.05.036. PubMed PMID: 34171319; PMCID: PMC8336675.

8. Keyes BE, Liu S, Asare A, Naik S, Levorse J, Polak L, Lu CP, Nikolova M, Pasolli HA, Fuchs E. Impaired Epidermal to Dendritic T Cell Signaling Slows Wound Repair in Aged Skin. Cell. 2016;167(5):1323–38 e14. doi: 10.1016/j.cell.2016.10.052. PubMed PMID: 27863246; PMCID: PMC5364946.

9. Mempel TR, Henrickson SE, Von Andrian UH. T-cell priming by dendritic cells in lymph nodes occurs in three distinct phases. Nature. 2004;427(6970):154–9. Epub 2004/01/09. doi: 10.1038/nature02238. PubMed PMID: 14712275.

10. Okada T, Miller MJ, Parker I, Krummel MF, Neighbors M, Hartley SB, O’Garra A, Cahalan MD, Cyster JG. Antigen-engaged B cells undergo chemotaxis toward the T zone and form motile conjugates with helper T cells. PLoS Biol. 2005;3(6):e150. Epub 2005/04/29. doi: 10.1371/journal.pbio.0030150. PubMed PMID: 15857154.

11. Chudnovskiy A, Pasqual G, Victora GD. Studying interactions between dendritic cells and T cells in vivo. Curr Opin Immunol. 2019;58:24–30. Epub 2019/03/19. doi: 10.1016/j.coi.2019.02.002. PubMed PMID: 30884422; PMCID: PMC6927575.

12. Moses L, Pachter L. Museum of spatial transcriptomics. Nat Methods. 2022;19(5):534–46. Epub 20220310. doi: 10.1038/s41592-022-01409-2. PubMed PMID: 35273392.

13. Moffitt JR, Lundberg E, Heyn H. The emerging landscape of spatial profiling technologies. Nat Rev Genet. 2022. Epub 2022/07/21. doi: 10.1038/s41576-022-00515-3. PubMed PMID: 35859028.

14. Maynard KR, Collado-Torres L, Weber LM, Uytingco C, Barry BK, Williams SR, Catallini JL, 2nd, Tran MN, Besich Z, Tippani M, Chew J, Yin Y, Kleinman JE, Hyde TM, Rao N, Hicks SC, Martinowich K, Jaffe AE. Transcriptome-scale spatial gene expression in the human dorsolateral prefrontal cortex. Nat Neurosci. 2021;24(3):425–36. Epub 20210208. doi: 10.1038/s41593-020-00787-0. PubMed PMID: 33558695; PMCID: PMC8095368.

15. Rodriques SG, Stickels RR, Goeva A, Martin CA, Murray E, Vanderburg CR, Welch J, Chen LM, Chen F, Macosko EZ. Slide-seq: A scalable technology for measuring genome-wide expression at high spatial resolution. Science. 2019;363(6434):1463–7. Epub 20190328. doi: 10.1126/science.aaw1219. PubMed PMID: 30923225; PMCID: PMC6927209.

16. Liu S, Iorgulescu JB, Li S, Borji M, Barrera-Lopez IA, Shanmugam V, Lyu H, Morriss JW, Garcia ZN, Murray E, Reardon DA, Yoon CH, Braun DA, Livak KJ, Wu CJ, Chen F. Spatial maps of T cell receptors and transcriptomes reveal distinct immune niches and interactions in the adaptive immune response. Immunity. 2022;55(10):1940–52 e5. doi: 10.1016/j.immuni.2022.09.002. PubMed PMID: 36223726; PMCID: PMC9745674.

17. Andrews N, Serviss JT, Geyer N, Andersson AB, Dzwonkowska E, Sutevski I, Heijboer R, Baryawno N, Gerling M, Enge M. An unsupervised method for physical cell interaction profiling of complex tissues. Nat Methods. 2021;18(8):912–20. Epub 2021/07/14. doi: 10.1038/s41592-021-01196-2. PubMed PMID: 34253926.

18. Armingol E, Officer A, Harismendy O, Lewis NE. Deciphering cell-cell interactions and communication from gene expression. Nat Rev Genet. 2021;22(2):71–88. Epub 2020/11/11. doi: 10.1038/s41576-020-00292-x. PubMed PMID: 33168968; PMCID: PMC7649713.

19. Raredon MSB, Adams TS, Suhail Y, Schupp JC, Poli S, Neumark N, Leiby KL, Greaney AM, Yuan Y, Horien C, Linderman G, Engler AJ, Boffa DJ, Kluger Y, Rosas IO, Levchenko A, Kaminski N, Niklason LE. Single-cell connectomic analysis of adult mammalian lungs. Sci Adv. 2019;5(12):eaaw3851. Epub 2019/12/17. doi: 10.1126/sciadv.aaw3851. PubMed PMID: 31840053; PMCID: PMC6892628.

20. Ramilowski JA, Goldberg T, Harshbarger J, Kloppmann E, Lizio M, Satagopam VP, Itoh M, Kawaji H, Carninci P, Rost B, Forrest AR. A draft network of ligand-receptor-mediated multicellular signalling in human. Nat Commun. 2015;6:7866. Epub 2015/07/23. doi: 10.1038/ncomms8866. PubMed PMID: 26198319; PMCID: PMC4525178.

21. Shilts J, Severin Y, Galaway F, Muller-Sienerth N, Chong ZS, Pritchard S, Teichmann S, Vento-Tormo R, Snijder B, Wright GJ. A physical wiring diagram for the human immune system. Nature. 2022;608(7922):397–404. Epub 2022/08/04. doi: 10.1038/s41586-022-05028-x. PubMed PMID: 35922511.

22. Efremova M, Vento-Tormo M, Teichmann SA, Vento-Tormo R. CellPhoneDB: inferring cell-cell communication from combined expression of multi-subunit ligand-receptor complexes. Nature protocols. 2020;15(4):1484–506. Epub 2020/02/28. doi: 10.1038/s41596-020-0292-x. PubMed PMID: 32103204.

23. Liu DS, Loh KH, Lam SS, White KA, Ting AY. Imaging trans-cellular neurexin-neuroligin interactions by enzymatic probe ligation. PLoS One. 2013;8(2):e52823. Epub 2013/03/05. doi: 10.1371/journal.pone.0052823. PubMed PMID: 23457442; PMCID: PMC3573046.

24. Pasqual G, Chudnovskiy A, Tas JMJ, Agudelo M, Schweitzer LD, Cui A, Hacohen N, Victora GD. Monitoring T cell-dendritic cell interactions in vivo by intercellular enzymatic labelling. Nature. 2018;553(7689):496–500. doi: 10.1038/nature25442. PubMed PMID: 29342141.

25. Ombrato L, Nolan E, Kurelac I, Mavousian A, Bridgeman VL, Heinze I, Chakravarty P, Horswell S, Gonzalez-Gualda E, Matacchione G, Weston A, Kirkpatrick J, Husain E, Speirs V, Collinson L, Ori A, Lee JH, Malanchi I. Metastatic-niche labelling reveals parenchymal cells with stem features. Nature. 2019;572(7771):603–8. Epub 20190828. doi: 10.1038/s41586-019-1487-6. PubMed PMID: 31462798; PMCID: PMC6797499.

26. Zhang S, Zhao H, Liu Z, Liu K, Zhu H, Pu W, He L, Wang RA, Zhou B. Monitoring of cell-cell communication and contact history in mammals. Science. 2022;378(6623):eabo5503. Epub 20221202. doi: 10.1126/science.abo5503. PubMed PMID: 36454848.

27. Bechtel TJ, Reyes-Robles T, Fadeyi OO, Oslund RC. Strategies for monitoring cell-cell interactions. Nat Chem Biol. 2021;17(6):641–52. Epub 20210525. doi: 10.1038/s41589-021-00790-x. PubMed PMID: 34035514.

28. Mazmanian SK, Liu G, Ton-That H, Schneewind O. Staphylococcus aureus sortase, an enzyme that anchors surface proteins to the cell wall. Science. 1999;285(5428):760–3. PubMed PMID: 10427003.

29. Guimaraes CP, Witte MD, Theile CS, Bozkurt G, Kundrat L, Blom AE, Ploegh HL. Site-specific C-terminal and internal loop labeling of proteins using sortase-mediated reactions. Nature protocols. 2013;8(9):1787–99. doi: 10.1038/nprot.2013.101. PubMed PMID: 23989673; PMCID: PMC3943461.

30. Dorr BM, Ham HO, An C, Chaikof EL, Liu DR. Reprogramming the specificity of sortase enzymes. Proc Natl Acad Sci U S A. 2014;111(37):13343–8. doi: 10.1073/pnas.1411179111. PubMed PMID: 25187567; PMCID: PMC4169943.

31. Dustin ML, Depoil D. New insights into the T cell synapse from single molecule techniques. Nat Rev Immunol. 2011;11(10):672–84. Epub 2011/09/10. doi: 10.1038/nri3066. PubMed PMID: 21904389; PMCID: PMC3889200.

32. Madisen L, Zwingman TA, Sunkin SM, Oh SW, Zariwala HA, Gu H, Ng LL, Palmiter RD, Hawrylycz MJ, Jones AR, Lein ES, Zeng H. A robust and high-throughput Cre reporting and characterization system for the whole mouse brain. Nature neuroscience. 2010;13(1):133–40. Epub 2009/12/22. doi: 10.1038/nn.2467. PubMed PMID: 20023653; PMCID: 2840225.

33. Stoll S, Delon J, Brotz TM, Germain RN. Dynamic imaging of T cell-dendritic cell interactions in lymph nodes. Science. 2002;296(5574):1873–6. Epub 2002/06/08. doi: 10.1126/science.1071065. PubMed PMID: 12052961.

34. Frederico B, Martins I, Chapela D, Gasparrini F, Chakravarty P, Ackels T, Piot C, Almeida B, Carvalho J, Ciccarelli A, Peddie CJ, Rogers N, Briscoe J, Guillemot F, Schaefer AT, Saude L, Reis ESC. DNGR-1-tracing marks an ependymal cell subset with damage-responsive neural stem cell potential. Dev Cell. 2022;57(16):1957–75 e9. doi: 10.1016/j.devcel.2022.07.012. PubMed PMID: 35998585; PMCID: PMC9616800.

35. Rubtsov YP, Niec RE, Josefowicz S, Li L, Darce J, Mathis D, Benoist C, Rudensky AY. Stability of the regulatory T cell lineage in vivo. Science. 2010;329(5999):1667–71. Epub 2010/10/12. doi: 10.1126/science.1191996. PubMed PMID: 20929851.

36. Aghajani K, Keerthivasan S, Yu Y, Gounari F. Generation of CD4CreER(T(2)) transgenic mice to study development of peripheral CD4-T-cells. Genesis. 2012;50(12):908–13. Epub 20120912. doi: 10.1002/dvg.22052. PubMed PMID: 22887772; PMCID: PMC3535561.

37. Shulman Z, Gitlin AD, Targ S, Jankovic M, Pasqual G, Nussenzweig MC, Victora GD. T follicular helper cell dynamics in germinal centers. Science. 2013;341(6146):673–7. Epub 2013/07/28. doi: 10.1126/science.1241680. PubMed PMID: 23887872; PMCID: 3941467.

38. Dogan I, Bertocci B, Vilmont V, Delbos F, Megret J, Storck S, Reynaud CA, Weill JC. Multiple layers of B cell memory with different effector functions. Nat Immunol. 2009;10(12):1292–9. Epub 2009/10/27. doi: 10.1038/ni.1814. PubMed PMID: 19855380.

39. McDonald BD, Jabri B, Bendelac A. Diverse developmental pathways of intestinal intraepithelial lymphocytes. Nat Rev Immunol. 2018;18(8):514–25. Epub 2018/05/03. doi: 10.1038/s41577-018-0013-7. PubMed PMID: 29717233; PMCID: PMC6063796.

40. el Marjou F, Janssen KP, Chang BH, Li M, Hindie V, Chan L, Louvard D, Chambon P, Metzger D, Robine S. Tissue-specific and inducible Cre-mediated recombination in the gut epithelium. Genesis. 2004;39(3):186–93. Epub 2004/07/30. doi: 10.1002/gene.20042. PubMed PMID: 15282745.

41. Mucida D, Husain MM, Muroi S, van Wijk F, Shinnakasu R, Naoe Y, Reis BS, Huang Y, Lambolez F, Docherty M, Attinger A, Shui JW, Kim G, Lena CJ, Sakaguchi S, Miyamoto C, Wang P, Atarashi K, Park Y, Nakayama T, Honda K, Ellmeier W, Kronenberg M, Taniuchi I, Cheroutre H. Transcriptional reprogramming of mature CD4(+) helper T cells generates distinct MHC class II-restricted cytotoxic T lymphocytes. Nat Immunol. 2013;14(3):281–9. Epub 2013/01/22. doi: 10.1038/ni.2523. PubMed PMID: 23334788; PMCID: PMC3581083.

42. Bilate AM, London M, Castro TBR, Mesin L, Bortolatto J, Kongthong S, Harnagel A, Victora GD, Mucida D. T Cell Receptor Is Required for Differentiation, but Not Maintenance, of Intestinal CD4(+) Intraepithelial Lymphocytes. Immunity. 2020;53(5):1001–14 e20. Epub 2020/10/07. doi: 10.1016/j.immuni.2020.09.003. PubMed PMID: 33022229; PMCID: PMC7677182.

43. Cepek KL, Shaw SK, Parker CM, Russell GJ, Morrow JS, Rimm DL, Brenner MB. Adhesion between epithelial cells and T lymphocytes mediated by E-cadherin and the alpha E beta 7 integrin. Nature. 1994;372(6502):190–3. doi: 10.1038/372190a0. PubMed PMID: 7969453.

44. Zen K, Liu Y, McCall IC, Wu T, Lee W, Babbin BA, Nusrat A, Parkos CA. Neutrophil migration across tight junctions is mediated by adhesive interactions between epithelial coxsackie and adenovirus receptor and a junctional adhesion molecule-like protein on neutrophils. Mol Biol Cell. 2005;16(6):2694–703. Epub 20050330. doi: 10.1091/mbc.e05-01-0036. PubMed PMID: 15800062; PMCID: PMC1142417.

45. Cohen CJ, Shieh JT, Pickles RJ, Okegawa T, Hsieh JT, Bergelson JM. The coxsackievirus and adenovirus receptor is a transmembrane component of the tight junction. Proc Natl Acad Sci U S A. 2001;98(26):15191–6. Epub 20011204. doi: 10.1073/pnas.261452898. PubMed PMID: 11734628; PMCID: PMC65005.

46. Pazirandeh A, Sultana T, Mirza M, Rozell B, Hultenby K, Wallis K, Vennstrom B, Davis B, Arner A, Heuchel R, Lohr M, Philipson L, Sollerbrant K. Multiple phenotypes in adult mice following inactivation of the Coxsackievirus and Adenovirus Receptor (Car) gene. PLoS One. 2011;6(6):e20203. Epub 20110603. doi: 10.1371/journal.pone.0020203. PubMed PMID: 21674029; PMCID: PMC3108585.

47. Subramanian A, Tamayo P, Mootha VK, Mukherjee S, Ebert BL, Gillette MA, Paulovich A, Pomeroy SL, Golub TR, Lander ES, Mesirov JP. Gene set enrichment analysis: a knowledge-based approach for interpreting genome-wide expression profiles. Proc Natl Acad Sci U S A. 2005;102(43):15545–50. Epub 2005/10/04. doi: 10.1073/pnas.0506580102. PubMed PMID: 16199517; PMCID: 1239896.

48. Stevens AJ, Harris AR, Gerdts J, Kim KH, Trentesaux C, Ramirez JT, McKeithan WL, Fattahi F, Klein OD, Fletcher DA, Lim WA. Programming Multicellular Assembly with Synthetic Cell Adhesion Molecules. Nature. 2022. Epub 20221212. doi: 10.1038/s41586-022-05622-z. PubMed PMID: 36509107.

49. Giladi A, Cohen M, Medaglia C, Baran Y, Li B, Zada M, Bost P, Blecher-Gonen R, Salame TM, Mayer JU, David E, Ronchese F, Tanay A, Amit I. Dissecting cellular crosstalk by sequencing physically interacting cells. Nat Biotechnol. 2020;38(5):629–37. Epub 2020/03/11. doi: 10.1038/s41587-020-0442-2. PubMed PMID: 32152598.

50. Sujino T, London M, Hoytema van Konijnenburg DP, Rendon T, Buch T, Silva HM, Lafaille JJ, Reis BS, Mucida D. Tissue adaptation of regulatory and intraepithelial CD4(+) T cells controls gut inflammation. Science. 2016;352(6293):1581–6. doi: 10.1126/science.aaf3892. PubMed PMID: 27256884; PMCID: PMC4968079.

51. Engels B, Cam H, Schuler T, Indraccolo S, Gladow M, Baum C, Blankenstein T, Uckert W. Retroviral vectors for high-level transgene expression in T lymphocytes. Hum Gene Ther. 2003;14(12):1155–68. doi: 10.1089/104303403322167993. PubMed PMID: 12908967.

52. Kim JH, Lee SR, Li LH, Park HJ, Park JH, Lee KY, Kim MK, Shin BA, Choi SY. High cleavage efficiency of a 2A peptide derived from porcine teschovirus-1 in human cell lines, zebrafish and mice. PLoS One. 2011;6(4):e18556. Epub 20110429. doi: 10.1371/journal.pone.0018556. PubMed PMID: 21602908; PMCID: PMC3084703.

53. Argos P. An investigation of oligopeptides linking domains in protein tertiary structures and possible candidates for general gene fusion. J Mol Biol. 1990;211(4):943–58. doi: 10.1016/0022-2836(90)90085-Z. PubMed PMID: 2313701.

54. Lee PP, Fitzpatrick DR, Beard C, Jessup HK, Lehar S, Makar KW, Perez-Melgosa M, Sweetser MT, Schlissel MS, Nguyen S, Cherry SR, Tsai JH, Tucker SM, Weaver WM, Kelso A, Jaenisch R, Wilson CB. A critical role for Dnmt1 and DNA methylation in T cell development, function, and survival. Immunity. 2001;15(5):763–74. Epub 2001/12/01. doi: 10.1016/s1074-7613(01)00227-8. PubMed PMID: 11728338.

55. Schraml BU, van Blijswijk J, Zelenay S, Whitney PG, Filby A, Acton SE, Rogers NC, Moncaut N, Carvajal JJ, Reis e Sousa C. Genetic tracing via DNGR-1 expression history defines dendritic cells as a hematopoietic lineage. Cell. 2013;154(4):843–58. doi: 10.1016/j.cell.2013.07.014. PubMed PMID: 23953115.

56. Shinnakasu R, Inoue T, Kometani K, Moriyama S, Adachi Y, Nakayama M, Takahashi Y, Fukuyama H, Okada T, Kurosaki T. Regulated selection of germinal-center cells into the memory B cell compartment. Nat Immunol. 2016;17(7):861–9. Epub 2016/05/10. doi: 10.1038/ni.3460. PubMed PMID: 27158841.

57. Barnden MJ, Allison J, Heath WR, Carbone FR. Defective TCR expression in transgenic mice constructed using cDNA-based alpha- and beta-chain genes under the control of heterologous regulatory elements. Immunology and cell biology. 1998;76(1):34–40. Epub 1998/04/29. doi: 10.1046/j.1440-1711.1998.00709.x. PubMed PMID: 9553774.

58. Danciu C, Falamas A, Dehelean C, Soica C, Radeke H, Barbu-Tudoran L, Bojin F, Pinzaru SC, Munteanu MF. A characterization of four B16 murine melanoma cell sublines molecular fingerprint and proliferation behavior. Cancer Cell Int. 2013;13:75. Epub 20130726. doi: 10.1186/1475-2867-13-75. PubMed PMID: 23890195; PMCID: PMC3750233.

59. Pasqual G, Angelini A, Victora GD. Triggering positive selection of germinal center B cells by antigen targeting to DEC-205. Methods in molecular biology. 2015;1291:125–34. Epub 2015/04/04. doi: 10.1007/978-1-4939-2498-1_10. PubMed PMID: 25836306.

60. Bilate AM, Bousbaine D, Mesin L, Agudelo M, Leube J, Kratzert A, Dougan SK, Victora GD, Ploegh HL. Tissue-specific emergence of regulatory and intraepithelial T cells from a clonal T cell precursor. Sci Immunol. 2016;1(2):eaaf7471. doi: 10.1126/sciimmunol.aaf7471. PubMed PMID: 28783695.

61. Wolf FA, Angerer P, Theis FJ. SCANPY: large-scale single-cell gene expression data analysis. Genome biology. 2018;19(1):15. Epub 2018/02/08. doi: 10.1186/s13059-017-1382-0. PubMed PMID: 29409532; PMCID: PMC5802054.

62. Sturm G, Szabo T, Fotakis G, Haider M, Rieder D, Trajanoski Z, Finotello F. Scirpy: a Scanpy extension for analyzing single-cell T-cell receptor-sequencing data. Bioinformatics. 2020;36(18):4817–8. doi: 10.1093/bioinformatics/btaa611. PubMed PMID: 32614448; PMCID: PMC7751015.

63. Franzen O, Gan LM, Bjorkegren JLM. PanglaoDB: a web server for exploration of mouse and human single-cell RNA sequencing data. Database (Oxford). 2019;2019. doi: 10.1093/database/baz046. PubMed PMID: 30951143; PMCID: PMC6450036.

64. Setty M, Tadmor MD, Reich-Zeliger S, Angel O, Salame TM, Kathail P, Choi K, Bendall S, Friedman N, Pe’er D. Wishbone identifies bifurcating developmental trajectories from single-cell data. Nat Biotechnol. 2016;34(6):637–45. Epub 20160502. doi: 10.1038/nbt.3569. PubMed PMID: 27136076; PMCID: PMC4900897.

65. Storey JD, Tibshirani R. Statistical significance for genomewide studies. Proc Natl Acad Sci U S A. 2003;100(16):9440–5. Epub 20030725. doi: 10.1073/pnas.1530509100. PubMed PMID: 12883005; PMCID: PMC170937.

66. Jumper J, Evans R, Pritzel A, Green T, Figurnov M, Ronneberger O, Tunyasuvunakool K, Bates R, Zidek A, Potapenko A, Bridgland A, Meyer C, Kohl SAA, Ballard AJ, Cowie A, Romera-Paredes B, Nikolov S, Jain R, Adler J, Back T, Petersen S, Reiman D, Clancy E, Zielinski M, Steinegger M, Pacholska M, Berghammer T, Bodenstein S, Silver D, Vinyals O, Senior AW, Kavukcuoglu K, Kohli P, Hassabis D. Highly accurate protein structure prediction with AlphaFold. Nature. 2021;596(7873):583–9. Epub 20210715. doi: 10.1038/s41586-021-03819-2. PubMed PMID: 34265844; PMCID: PMC8371605.

67. Emsley P, Cowtan K. Coot: model-building tools for molecular graphics. Acta Crystallogr D Biol Crystallogr. 2004;60(Pt 12 Pt 1):2126–32. Epub 20041126. doi: 10.1107/S0907444904019158. PubMed PMID: 15572765.

68. Ko J, Park H, Heo L, Seok C. GalaxyWEB server for protein structure prediction and refinement. Nucleic Acids Res. 2012;40(Web Server issue):W294–7. Epub 20120530. doi: 10.1093/nar/gks493. PubMed PMID: 22649060; PMCID: PMC3394311.

69. Jo S, Kim T, Iyer VG, Im W. CHARMM-GUI: a web-based graphical user interface for CHARMM. J Comput Chem. 2008;29(11):1859–65. doi: 10.1002/jcc.20945. PubMed PMID: 18351591.

70. Paulick MG, Bertozzi CR. The glycosylphosphatidylinositol anchor: a complex membrane-anchoring structure for proteins. Biochemistry. 2008;47(27):6991–7000. Epub 20080617. doi: 10.1021/bi8006324. PubMed PMID: 18557633; PMCID: PMC2663890.

71. Goddard TD, Huang CC, Meng EC, Pettersen EF, Couch GS, Morris JH, Ferrin TE. UCSF ChimeraX: Meeting modern challenges in visualization and analysis. Protein Sci. 2018;27(1):14–25. Epub 20170906. doi: 10.1002/pro.3235. PubMed PMID: 28710774; PMCID: PMC5734306.

72. Schrödinger LLC, DeLano W. PyMOL v.2.4.02020.

73. Butler A, Hoffman P, Smibert P, Papalexi E, Satija R. Integrating single-cell transcriptomic data across different conditions, technologies, and species. Nat Biotechnol. 2018;36(5):411–20. Epub 2018/04/03. doi: 10.1038/nbt.4096. PubMed PMID: 29608179; PMCID: PMC6700744.

